# Water-mediated productivity dynamics in shifting coral reef communities

**DOI:** 10.64898/2026.04.17.719123

**Authors:** Jana Vetter, Kara E. Engelhardt-Stolz, André Dietzmann, Franziska Wöhrmann-Zipf, Maren Ziegler

## Abstract

Mass mortality of reef-building stony corals has driven widespread community shifts towards reefs dominated by soft corals and macroalgae. Although physical competition for space between these organisms plays an important role, non-contact water-mediated interactions have been proposed to modulate organismal performance and community functioning, yet their independent effects remain poorly resolved. Here, we experimentally tested the hypothesis that water-mediated interactions generate non-additive effects on community productivity, altering ecosystem functioning during phase shifts. Using two controlled incubation experiments with representative stony corals, soft corals, and macroalgae, we compared monoculture baseline productivity with mixed assemblages across a gradient of biomass ratios mimicking phase shift scenarios. We found that reductions in stony coral biomass led to community-level declines in photosynthesis and calcification that exceeded expectations based on monocultures, indicating emergent negative effects of community restructuring. However, these effects were strongly species-dependent, with some assemblages showing only minor deviations from expectations, whereas others exhibited pronounced productivity losses. At the species level, both stony corals reduced photosynthetic efficiency in mixed assemblages, while soft corals maintained efficiency across treatments. Macroalgal responses diverged, with one species exhibiting reduced and another increased photosynthetic efficiency in mixed communities. These species-specific physiological responses scaled up to explain community-level deviations from expected productivity, suggesting that gains in productivity by certain taxa can partially offset, but not fully compensate for, losses in coral-driven functions such as calcification. Together, our findings indicate that sublethal, water-mediated interactions can reorganize holobiont functioning and lead to changes in ecosystem productivity, independent of direct physical competition. By altering community-wide energy acquisition and carbonate production, such interactions may reinforce feedback loops that accelerate ecosystem phase shifts. We argue that incorporating water-mediated interaction effects into ecological theory and ecosystem models is essential for predicting the stability and recovery potential of coral reefs and other transitioning ecosystems under climate change.

## Introduction

Mass extinction of stony corals has led to phase shifts of coral reefs to alternative stable states globally in the last decades (Norström et al., 2009). When stony corals die off, opportunistic species, such as macroalgae and soft corals, immediately start to grow over the bare substrate, occupying the space needed for larval settlement and coral recovery. This can lead to a negative feedback loop, resulting in a ‘phase shift’ from reefs dominated by stony coral biomass to those dominated by macroalgae or soft coral biomass (Done, T.J., 1992). As a consequence, coral reefs ultimately undergo a reduction in their three-dimensional structure (Done, T.J., 1992; Norström et al., 2009), which triggers drastic ecosystem changes, with negative impacts on ecosystem services and cascading effects on adjacent ecosystems (Reaka-Kudla, 1997). While impacts vary by region and reef, alternative stable states usually result in lower biodiversity, lower community calcification, higher net community photosynthesis, along with other changes in carbon flux (Done, T.J., 1992; Roth et al., 2021). Multiple examples of such phase shift events are documented worldwide, spanning from the Caribbean (Mumby, 2009), the Indian Ocean (Graham et al., 2015; McManus & Polsenberg, 2004), the Red Sea (Anton et al., 2020; Riegl & Piller, 1999), to the Pacific (Roff et al., 2015).

Phase shifts typically start with a pulse-stress event, such as a disease outbreak or coral bleaching, that initiate a negative feedback loop which is further driven by additional anthropogenic stressors such as overfishing of herbivores and water pollution (McManus & Polsenberg, 2004). Even when the stressors are reduced, negative feedback loops may still shift the coral-dominated reef to an alternative stable state of soft coral- or macroalgae-dominance. This phenomenon results from intricate biological interactions among the competing benthic organisms, the mechanisms of which remain insufficiently understood. A comprehensive understanding of these feedback loops is crucial for devising better management strategies to prevent or reverse phase shifts in the future.

Both physical and water-mediated interactions among competing benthic organisms can impact one another. Physical contact can impair coral health resulting in further decline of stony coral biomass (Clements & Hay, 2023; Thurber et al., 2012). Stony corals can also sense other organisms through water-mediated interactions, which can result in productivity alterations (Engelhardt et al., 2023). Known water-mediated interaction pathways between stony corals, soft corals, and macroalgae include secondary metabolites (Clements & Hay, 2023; Rodriguez et al., 2021) and microbial interactions (Briggs et al., 2021; Fong et al., 2020; Haas et al., 2013), which can reduce growth rate (Fu et al., 2023), polyp activity (Rodriguez et al., 2021), photosynthesis (Engelhardt et al., 2023), photosynthetic efficiency (Smith et al., 2006), symbiont density (Aceret et al., 1995; Engelhardt et al., 2023; Fu et al., 2023), and induce tissue necrosis (Aceret et al., 1995; Sammarco et al., 1983; Smith et al., 2006) and changes at the levels of gene expression in stony corals (Andrade Rodríguez, 2018; Fu et al., 2022). For instance, water-mediated interactions between the soft coral *Lobophytum* sp. and the stony coral *Porites* sp. resulted in a broad immune response of the stony coral, with upregulated genes that have previously been found to respond to contact with pathogenic bacteria, chemotoxic attacks in direct contact to macroalgae, and heat stress (Andrade Rodríguez, 2018). Therefore, water-mediated interactions without physical touch between organisms in the reef may be as stressful to stony corals as direct contact or heat stress (Andrade Rodríguez, 2018).

Stress responses manifest in the health of organisms that can be quantified as subtle physiological changes, including modulation of respiration and photosynthesis. These alterations can be a sign of resource divergence and an ‘indirect cost’ which can negatively affect organisms in the long-term (Huey et al., 2002). Engelhardt et al. (2023) demonstrated that short-term and long-term water-mediated interactions induce species-specific responses in stony coral productivity, which were mostly independent of the interaction partner. It is therefore crucial to understand these subtle responses of stony corals, as well as their competitors for space with soft corals and macroalgae, as this may give us essential clues into why some organisms and reef communities cope better than others during phase shifts.

This study aims to disentangle changes in community productivity and individual productivity through water-mediated interactions between stony corals, soft corals, and macroalgae in various biomass constellations as a model of phase shift scenarios on the reef. To this end, we conducted two fully controlled incubation experiments with two stony coral, two soft coral and two macroalga species. Our main objectives were: i) to quantify the baseline productivity of each species in monoculture; ii) to determine changes in community productivity in a ‘phase shift scenario’, by adjusting biomass ratios in polycultures of stony corals with either soft corals or macroalgae; iii) to determine changes in community productivity in a ‘degraded reef scenario’, with low or absent stony coral biomass in polycultures of stony corals, soft corals, and macroalgae or solely soft corals and macroalgae; iv) to identify the species likely responsible for changes in community productivity by assessing how individual species change productivity in response to other organisms.

## Materials & Methods

To explore the effect of water-mediated interactions on the productivity of benthic reef organisms, two incubation experiments with six sessile coral reef organisms from three organism groups (stony corals, soft corals, macroalgae) were performed. Photosynthesis, respiration, and calcification were measured as community productivity parameters in the first experiment. To determine how each species contributed to the observed productivity changes, a second experiment with the same species was conducted to measure changes in photosynthetic efficiency induced by the presence of other organisms for each species separately.

Both experiments were conducted in the *Ocean2100* aquarium facility and a temperature-controlled multi-point stirring incubator (Rades et al., 2022) of the Justus Liebig University Giessen, Germany. The species consisted of two stony corals (Tab. S1): *Montipora digitata* (Dana, 1846) & *Porites rus* (Forskal, 1775); two soft corals: *Xenia umbellata* (Lamarck, 1816) & *Sclerophytum wanannensis* (van Ofwegen & Benayahu, 2012; formerly *Sinularia wanannensis,* May, 1899); and two macroalgae: *Caulerpa brachypus* (Harvey, 1860) & *Peyssonnelia* sp. (Decaisne, 1841; Fig. 1a). The two stony coral species were selected based on their strong reaction to other stony corals species (Vetter et al., 2025). The soft coral and macroalgae species are strong competitors of stony corals. *Xenia* softcorals overgrow reefs rapidly and due to their resilience to warmer, more eutrophic water, are feared to outcompete stony corals in the future (Mezger et al., 2022; Norström et al., 2009). *Sclerophytum* softcorals and the macroalgae *Caulerpa* both produce allelopathic substances against competitors (Kase et al., 2020; Maida et al., 2001), with *Caulerpa* up- or downregulating its production during competition (Dumay et al., 2002). *Peyssonnelia* spp. are crustose algae, which is becoming more dominant on coral reefs by overgrowing stony corals, and through their resistance to climate change *Peyssonnelia* spp. pose a potential threat to coral reefs (Edmunds et al., 2023). The species were fragmented and kept separately at 26 °C and a salinity of 35, in 265-liter tanks during the healing/acclimation (12 weeks for the stony corals, 2 weeks for the soft corals & macroalgae) and experimental period. This ensured a naïve state of the organisms without exposure to stimuli from others. A 10:14 h light:dark cycle at ∼200 µmol photons m^-2^ s^-1^ was maintained. The stony corals and the macroalga *C*. *brachypus* were suspended from nylon lines and weighed down with a stainless-steel nut when necessary. The two soft corals and the macroalga *Peyssonnelia* sp. were glued to clay plugs and held on light grid plates.

**Figure 1.**
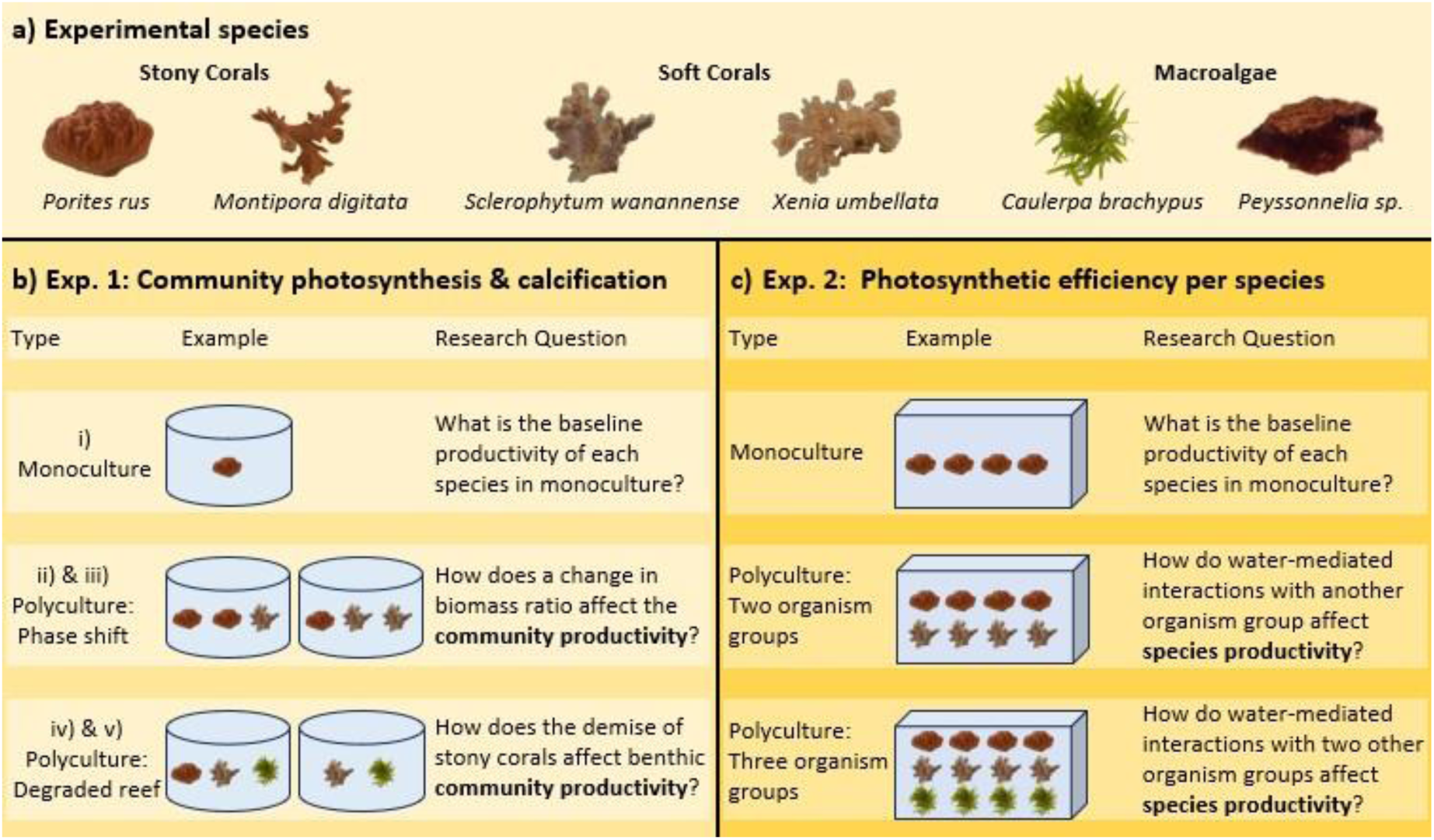
Overview of the experimental set-up and research questions. a) Experimental species: *P. rus* & *M. digitata* (stony corals), *S. wanannensis* sp. & *X. umbellata* (soft corals), *C. brachypus* and *Peyssonnelia* sp. (Macroalgae). b) Incubation types of the community productivity measurements. Productivity measurements from the monoculture incubations were used to test for changes caused by sessile reef neighbours in the polyculture incubations (i). The ‘phase shift’ polycultures were either composed of two stony coral fragments with one soft coral or one macroalga fragment (ii), or one stony coral fragment with two soft coral or two macroalga fragments (iii). The ‘degraded reef’ polycultures were either composed of one stony coral, one soft coral and one macroalga fragment (iv), or one soft coral and one macroalga fragment (v). c) Incubation types of the individual photosynthetic efficiency (YII) measurements. Measurements from the monoculture incubations were used to test for changes caused by sessile reef neighbours in two polyculture incubation types. The polycultures of two organism groups were composed of four stony coral fragments with four soft coral or four macroalgae fragments, each of the same species. The polycultures of the three organism groups were composed of four stony coral, four soft coral, and four macroalga fragments, each of the same species.

### Experiment 1

Changes in net photosynthesis, respiration, gross photosynthesis, and calcification through water-mediated interactions between sessile reef neighbours were investigated to quantify effects on community productivity. The productivity measurements were taken in airtight, 1-liter incubation jars over 52 days. During this time 559 incubations in the light and 559 incubations in the dark were conducted. Filtered (65 µm) aquarium water from an empty tank of the recirculation system of the holding tanks was used for the incubations. Nine replicate fragments per organism were used (n = 9). This meant three replicate fragments from three genotypes of each stony coral and nine replicate fragments from one genotype of each soft coral and macroalgae.

Five different types of incubations were performed (Fig. 1b). i) Monoculture incubations were conducted to determine the baseline productivity of each organism in their naïve state. To cover potential variations over the experimental period these incubations were performed at the start, middle, and the end of the experiment (n = 27). ii) Two stony coral fragments from the same species and genotype were incubated together with one soft coral or one macroalgae fragment, to investigate changes in productivity by sessile reef neighbours (n = 9). iii) To imitate a ‘phase shift’ one stony coral fragment was incubated with either two soft coral or two macroalgae fragments from the same species. This way the effect of reduced stony coral and increased soft coral or macroalgae biomass on their productivity could be investigated (n = 9). iv) One stony coral, one soft coral, and one macroalgae fragment were incubated together to investigate community productivity in a diverse reef setting (n = 9). v) To explore community productivity changes in a degraded reef without stony corals, one soft coral and one macroalgae fragment were incubated together (n = 9).

Incubation time was set to 60 minutes in the monoculture and 30 minutes in the polyculture incubations, to obtain measurable differences between the start and end of the incubations, while avoiding critical oxygen over or under saturation. Abiotic conditions were the same as in the holding tanks, with a light intensity between 195-205 µmol photons m^-2^ s^-1^, a flow rate of 6-7 cm s^-1^ and temperature of 26.0 ± 0.3 (SD) °C in the light and 25.9 ± 0.3 (SD) °C in the dark incubations. Organisms were incubated together with an approximate minimum distance of 3 cm. *C. brachypus* fragments grew rapidly and needed to be refragmented two times during the experiment to keep that distance to the other organisms. These fragments were given two days to heal before being used in the incubations again. All organisms were suspended in the incubation jars at a uniform depth to exclude overshadowing within the incubation jars and to prevent interferences with the magnetic stirrer at the bottom.

### Photosynthesis & respiration

Oxygen measurements were performed with an optical oxygen multiprobe (FDO 925, WTW Multi 3620 IDS, Weilheim, Germany). The difference in oxygen content between the start and end of the incubations was used to calculate net photosynthesis during the light and respiration during the dark incubations. Both productivity parameters were normalized with organism biomass (Ash free dry weight), water volume, and incubation time. To account for productivity from microorganism in the water, oxygen changes were corrected by subtracting the mean net photosynthesis/respiration of two seawater controls per day. Gross photosynthesis was calculated by adding up net photosynthesis and respiration values.

### Calcification

Calcification measurements were taken during the light incubations using the total alkalinity anomaly technique (Chisholm & Gattuso, 1991). For this, 50 ml water samples were collected at the beginning and end of each incubation, stored in the dark at room temperature (20 °C) and analysed within 5 h of collection. Total alkalinity (TA) of these samples was determined via potentiometric titration with 0.1 N HCL (SI Analytics TitroLine® 7000, Mainz, Germany) and a glass electrode (Pt1000 from SI Analytics™). The pH electrode was calibrated every day before the first measurements (slope from 98 to 99%) and had an accuracy of 0.002 pH. The Gran Alkalinity Approximation was used to calculate TA. Calcification rate was calculated as described by Schneider & Erez (2006) :

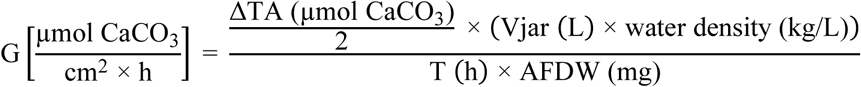

Where ΔTA is the difference between TA at the start and end of each incubation, Vjar is the water volume of the incubation jar (L), T is the incubation time (h), and AFDW represents the ash free dry weight of the incubated fragments (mg).

### Normalization to biomass (AFDW)

Ash free dry weight has been shown to be a good normalization method for organisms with different physiques (Pupier et al., 2018). Thus, to enable a productivity comparison among the three organism groups, the production values were normalized to ash free dry weight (AFDW, Tab. 1). Due to the length of the experiment, it was necessary to track the growth of the organisms with a nonlethal method, to ensure productivity values could be accurately normalized. Therefore, the wet weight of the soft coral and macroalgae fragments, as well as the buoyant weight of the stony coral fragments was measured at the start, middle and end of the experiment. The AFDW for these fragments was calculated based on a linear regression formular between wet weight and buoyant weight with AFDW. This formular was created beforehand for each species (Fig. S1). For this, fragments of each species were weighed and the stony coral tissue was separated from the skeleton using an air-brush. Subsequently the stony coral tissue as well as the soft coral and macroalgae fragments were dried in an oven at 60 °C for 72 h. After determination of the dry weight, these samples were incinerated in a muffle furnace for 5 ½ h at 500 °C. The resulting ash-weight was subtracted from the dry weight to calculate the AFDW.

**Table 1.**
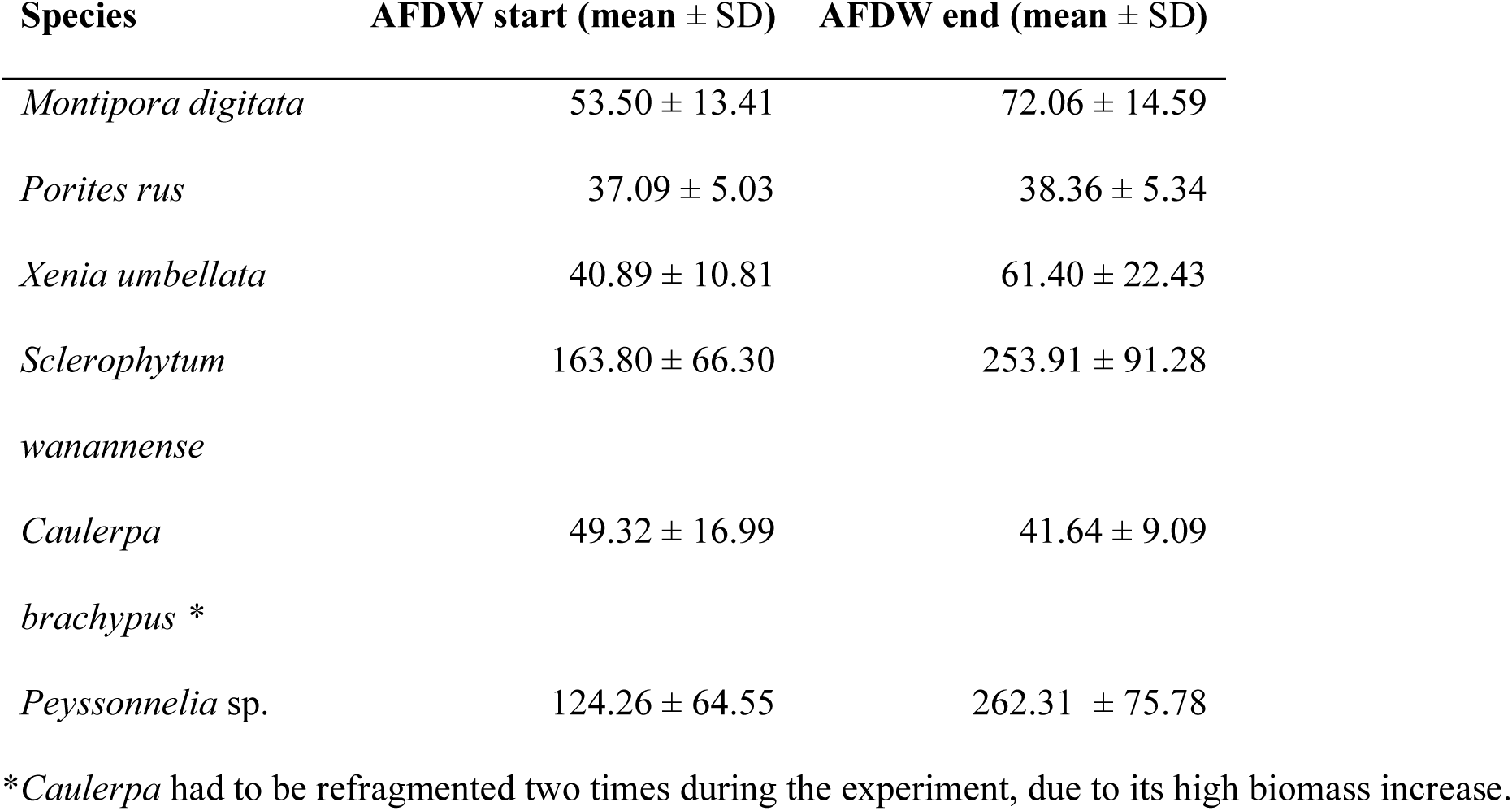
Ash free dry weight (AFDW) of all species at the beginning and the end of the experiment.

| Species | AFDW start (mean ± SD) | AFDW end (mean ± SD) |
| --- | --- | --- |
| *Montipora digitata* | 53.50 ± 13.41 | 72.06 ± 14.59 |
| *Porites rus* | 37.09 ± 5.03 | 38.36 ± 5.34 |
| *Xenia umbellata* | 40.89 ± 10.81 | 61.40 ± 22.43 |
| *Sclerophytum wanannense* | 163.80 ± 66.30 | 253.91 ± 91.28 |
| *Caulerpa brachypus* * | 49.32 ± 16.99 | 41.64 ± 9.09 |
| *Peyssonnelia* sp. | 124.26 ± 64.55 | 262.31 ± 75.78 |

\**Caulerpa* had to be refragmented two times during the experiment, due to its high biomass increase.

Species AFDW correlated positively with the productivity parameters (Fig. S2), except calcification with the non-calcifying species (*C. brachypus*, *X. umbellata* & *S. wanannensis*). This indicates a healthy productivity across all species and sizes. One exemption was *Peyssonnelia* sp. which showed no significant net photosynthesis increase with higher AFDW. The *Peyssonnelia* sp. fragments had various thicknesses, increasing/decreasing their AFDW without changing the light absorbing surface area, which might have led to a constant photosynthesis across the slightly varying fragment sizes.

### Experiment 2

To further disentangle water-mediated effects of experiment 1, changes in photosynthetic efficiency (YII) via pulse-amplitude modulation fluorometry (PAM) was investigated of individual organisms. The incubations took place in nine 30 x 15 cm aquaria which were filled with 7-liters of 65 µm filtered seawater from an empty tank of the recirculation system. Measurements took place over a total experimental period of 21 days. During this time 114 incubations were conducted. Twelve replicate fragments per organism were used (n = 12), i.e., four replicate fragments from three genotypes of each stony coral and twelve replicate fragments from one genotype of each soft coral and macroalgae. Fragmentation took place four weeks before the experiment to allow time to heal in monoculture tanks.

Three different types of incubations were performed (Fig. 1c). i) Monoculture incubations were conducted to determine the baseline YII values of each organism in their naïve state. To cover potential variations over the experimental period these incubations were performed at the start, middle and the end of the experiment. Three measurements were taken per fragment (n = 36). ii) Each stony coral was incubated together with one of the soft corals or macroalgae, to investigate changes in YII by sessile reef neighbours (n = 12). iii) One stony coral, one soft coral, and one macroalgae species were incubated together to investigate YII changes in a diverse reef setting (n = 12).

Measurements were taken after 30 minutes and 120 minutes of incubation. YII exhibited minimal variation between the two measurement points. Values measured after 120 minutes are shown in the results, except for the incubations in which *C. brachypus* and *M. digitata* were incubated together, for which the 30-minute measurements were used, as most of the 120-minute measurements were accidently deleted. Abiotic conditions were the same as before, except for the flow rate, which was between 0.3-1 cm s^-1^ in all incubations. Organisms were incubated together with an approximate minimum distance of 3 cm.

### Photosynthetic efficiency (YII)

Photosynthetic efficiency measurements were taken during light incubations with a ‘PAM-2500 Portable Chlorophyll Fluorometer’ (Heinz Walz GmbH, Germany) together with the ‘PamWin-3 System Control and Data Acquisition System’ software (V3.22d, Heinz Walz GmbH). Each fragment was measured three times at different spots after 30 minutes and 120 minutes of incubation. The mean of the triplicate measurements per time point was calculated for subsequent analysis and a cut-off of 0.1 SD was used to determine outliers among the three measurements. The settings during light impulses were the same for all species, with gain of 3 and Ft value of 200 ± 10. The distance to the organism surface was kept stable at 0.5 cm and an angle of 60° by attaching a small piece of silicon pipe to the fibreoptic.

### Statistical analysis

Data analysis was performed in R version 4.3.2 (R Core Team, 2023), using the tidyverse collection (Wickham et al., 2019). Three photosynthetic values, six calcification values and two PAM values were found to be outliers and removed from the dataset.

To evaluate differences in productivity among the six species based on their monoculture performance, a one-factorial ANOVA was conducted using the nlme package (Pinheiro et al., 2020). Post hoc comparisons were performed using Tukey’s test with Bonferroni correction, implemented via the multcomp package (Hothorn et al., 2008). To account for repeated measurements within individual fragments, ‘fragment ID’ was included as a random effect. The response variables ‘net photosynthesis’ and ‘gross photosynthesis’ were log-transformed to satisfy the assumptions of normality and homoscedasticity.

To assess the effect of alterations in stony coral biomass ratio on community productivity, paired t-test with Bonferroni correction were performed between the ‘Polyculture: Phase Shift’ incubations (see incubation types ii) and iii) of Fig. 1b). To further distinguish the productivity effect of lower stony coral biomass the difference between the measured community productivity of each polyculture assemblage and its expected productivity was determined. The expected productivity was calculated as the sum of the monoculture productivities of the corresponding fragments (Fig. 2). As each fragment was measured in monoculture three times (at the start, middle, and end of the experimental period), the values for intermediate days were estimated via point-point interpolation to generate daily monoculture productivity values used in the expected value calculation. To test for significant differences between expected and measured productivity, paired t-test with Bonferroni correction were performed. Wilcoxon signed rank tests were applied when parametric assumptions were not met. Results were visualized as the percentage change from expected productivity (Fig. 4e-h).

**Figure 2.**
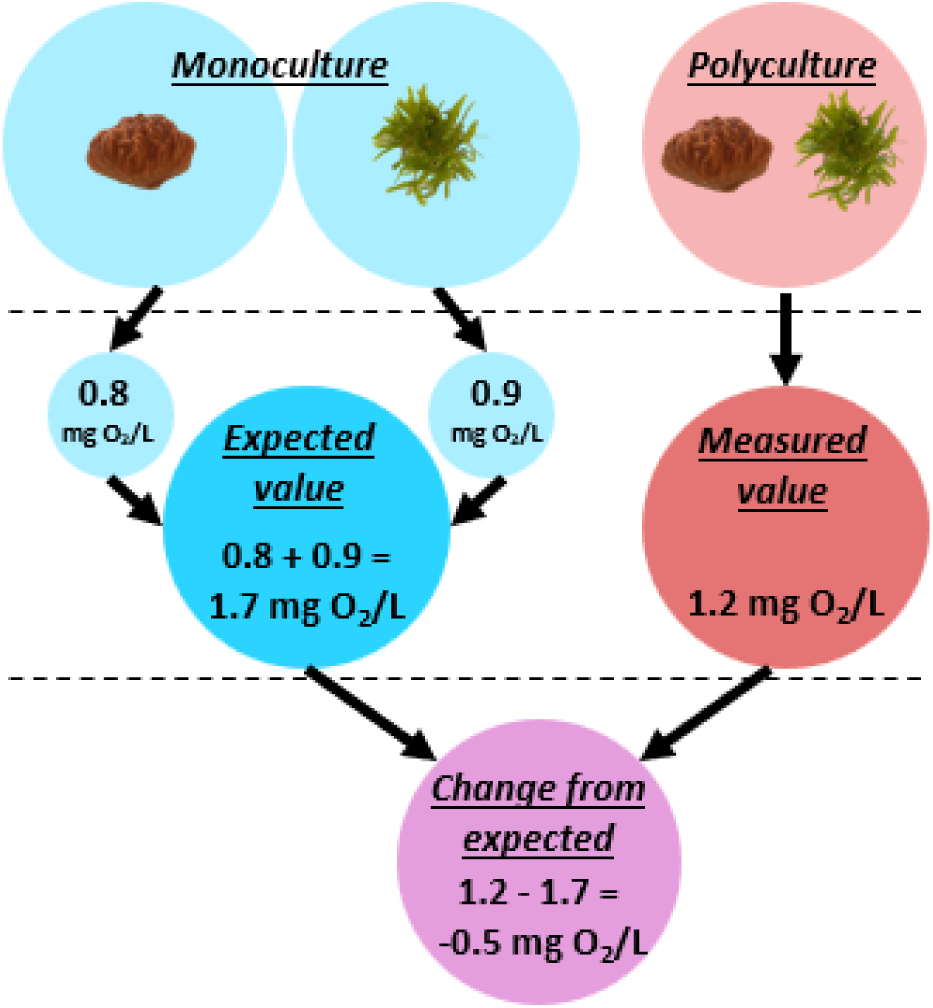
Calculation example expected value. The monoculture oxygen values of the same fragments used in the polyculture are summed to obtain the expected value. This expected value is then subtracted from the measured polyculture value to determine the change from expected.

To assess the effect of absent stony coral biomass on community productivity, a one-way ANOVA was conducted comparing polyculture incubations of the same soft coral and macroalgae species assemblage with and without the two stony coral species (see ‘Polyculture: Degraded reef’ incubation types iv) and v) of Fig. 1b). Post hoc comparisons were performed using Tukey’s test with Bonferroni correction. To account for repeated measurements within individual fragments, ‘fragment ID’ was included as a random effect. For incubations consisting of *S. wanannense* & *C. brachypus* the response variables ‘respiration’ and ‘gross photosynthesis’ were log-transformed to satisfy the assumptions of normality and homoscedasticity. The same was done for the response variable ‘gross photosynthesis’ for the incubations including *X. umbellata* & *Peyssonnelia* sp. To further distinguish the productivity effect of missing stony coral biomass, the difference between the measured community productivity of each polyculture assemblage and its expected productivity was determined via a Wilcoxon signed rank test, as parametric assumptions were not met. Results were visualized as the percentage change from expected productivity (Fig. 5d-f).

To evaluate the individual effect of other organism groups on stony coral productivity, a one-way ANOVA was conducted comparing photosynthetic efficiency measurements between monoculture incubations and corresponding polyculture incubations containing two and three organism groups (see incubation types in Fig. 1c). Post hoc comparisons were performed using Tukey’s test with Bonferroni correction. ‘fragment ID’ was included as a random effect to account for repeated measurements within individual fragments.

## Results

### Productivity differences of six sessile coral reef organisms

Net photosynthesis, respiration, gross photosynthesis, calcification, and photosynthetic efficiency were significantly different (p < 0.05) among the six coral reef organisms examined in monoculture (Fig. 3; Tab. S2). The photosynthetic productivity parameters showed a species-specific pattern with significant differences within as well as across organism groups, where *C. brachypus* was roughly twice as productive as the other species. Calcification was significantly higher in the two stony corals compared to the soft corals and macroalgae (p < 0.001), which did not calcify (Fig. 3d). For mean productivity values see Tab. S3.

**Figure 3.**
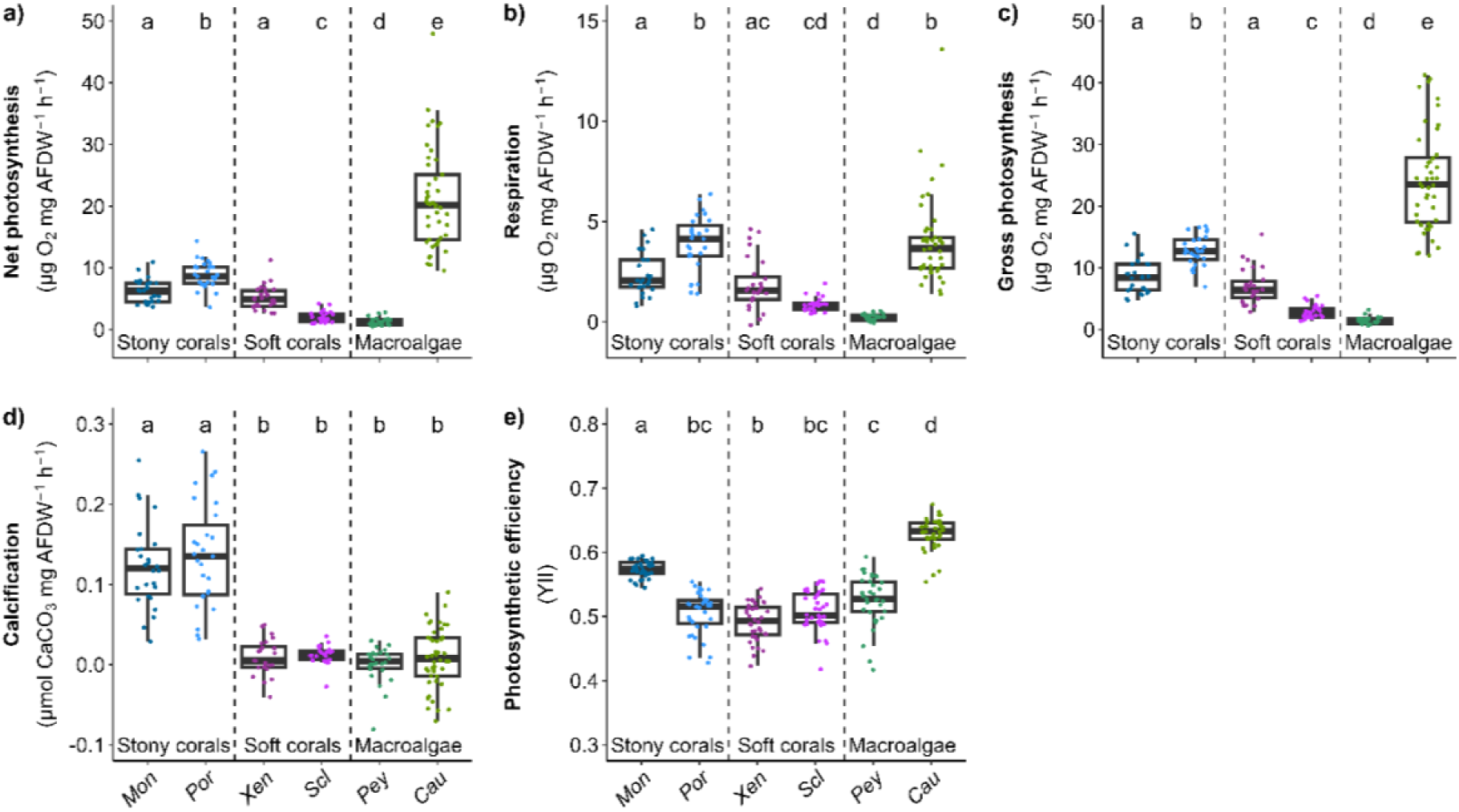
Productivity of six sessile coral reef organisms measured as net photosynthesis. (a), respiration (b), gross photosynthesis (c), calcification (d), and photosynthetic efficiency (e). Data based on monoculture incubations (Experiment 1 (a-d): n = 27, Experiment 2 (e): n = 108). Species abbreviations: Mon = *Montipora*, Por = *Porites*, Xen = *Xenia*, Scl = *Sclerophytum*, Cau = *Caulerpa,* Pey = *Peyssonnelia*. Boxplots are ordered by organism group, boxes indicate the first and third quartiles, and whiskers indicate ± 1.5 IQR. Letters indicate significant differences between organisms (p < 0.05, Tab. S2).

### Experiment 1

#### Effect of biomass ratios of competing organisms on community productivity

The comparison of community productivity between incubations with high stony coral biomass and those with higher soft coral or macroalgae biomass showed significant variations in productivity. An increase in soft coral and the subsequent decrease in stony coral biomass resulted in reduced community productivity (Fig. 4 a-d, Tab. S4). This aligns with the monoculture productivity findings whereby soft corals were less productive than stony corals (Fig. 3). Yet, the increase in soft corals led to a larger reduction in photosynthesis and respiration than anticipated based on the monoculture incubations (Fig. 4 e-h, Tab. S5), indicating a negative productivity impact on at least one of the interacting species. The largest impact was found for *M. digitata* incubations with an increase in *X. umbellata* biomass, resulting in a mean gross photosynthesis reduction from expected values of 38 % ± 11 (SD; Fig. 4g).

**Figure 4.**
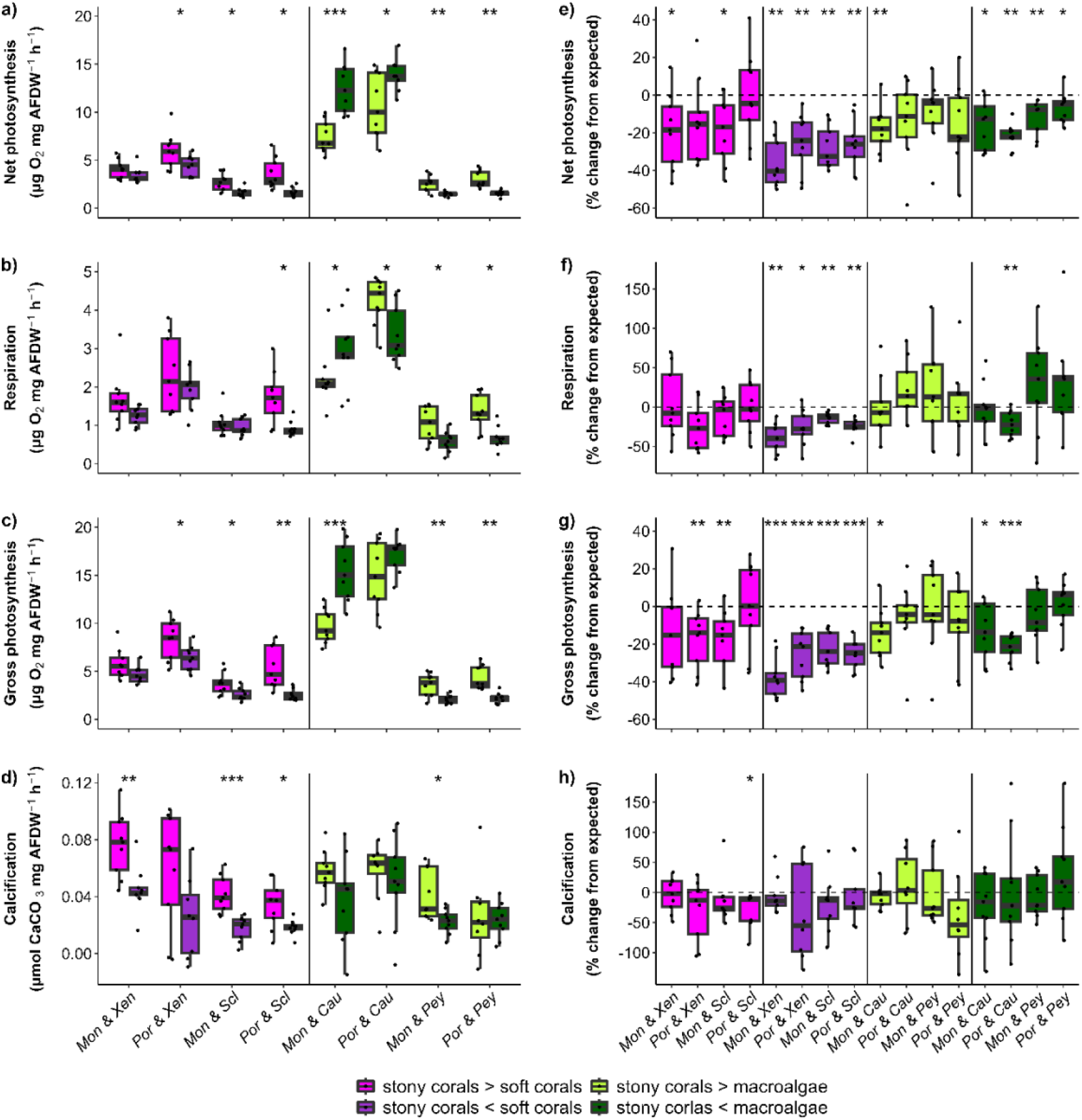
Shifts in biomass of reef neighbours affect community productivity. Community productivity is shown as net photosynthesis (a), respiration (b), gross photosynthesis (c), and calcification (d) in absolute values and as change from expected (in %) of net photosynthesis (e), respiration (f), gross photosynthesis (g), and calcification (h). Species abbreviations: Mon = *Montipora*, Por = *Porites*, Xen = *Xenia*, Scl = *Sclerophytum*, Cau = *Caulerpa,* Pey = *Peyssonnelia*. Boxes indicate the first and third quartiles, and whiskers indicate ± 1.5 IQR. Asterisks indicate significant difference between dominance incubations of the same species (a-d) or significant difference from expected (e-h; * p < 0.05, ** p < 0.01, *** p < 0.001; Tab. S4 & Tab. S5).

An increase in macroalgae and the simultaneous decrease in stony coral biomass resulted in contrasting effects, depending on the macroalgae species. Increased *C. brachypus* biomass enhanced community productivity, while an increase in *Peyssonnelia* sp. diminished overall productivity (Fig. 4 a-d). This again follows the same productivity pattern as observed in monocultures, where *C. brachypus* was among the most productive species, while *Peyssonnelia* sp. was one of the least productive (Fig. 3). However, compared to the expectations based on monoculture performances, increased biomass of both macroalgae mostly reduced community photosynthesis, with the greatest reduction seen in the incubations with increased *C. brachypus* biomass (Fig. 4 e-h). Specifically, elevated *C. brachypus* biomass drove the strongest decline in *P. rus* incubations, reducing gross photosynthesis by 22 % ± 7 (SD) relative to expectations. Interestingly, the communities with increased *Peyssonnelia* sp. biomass showed a positive non-significant trend in community productivity in all parameters except net photosynthesis. Notably, all incubations which encompassed a higher percentage of stony coral biomass than soft corals or macroalgae showed less pronounced changes from monoculture expectations. In other words, biomass ratios between stony corals and the other two benthic organism groups had a significant impact on community productivity, independent of the incubated species.

### Effect of stony coral loss on community productivity

To test the effect of stony coral cover on community productivity, incubations with one soft coral, one macroalgae, and one stony coral were compared to the same incubations without the stony corals. Community productivity with all three organisms was similar to that of incubations with only soft coral and macroalgae, except for communities containing *C. brachypus* and a soft coral, which were generally less productive without stony corals (Fig. 5 a-c, Tab. S6).

**Figure 5.**
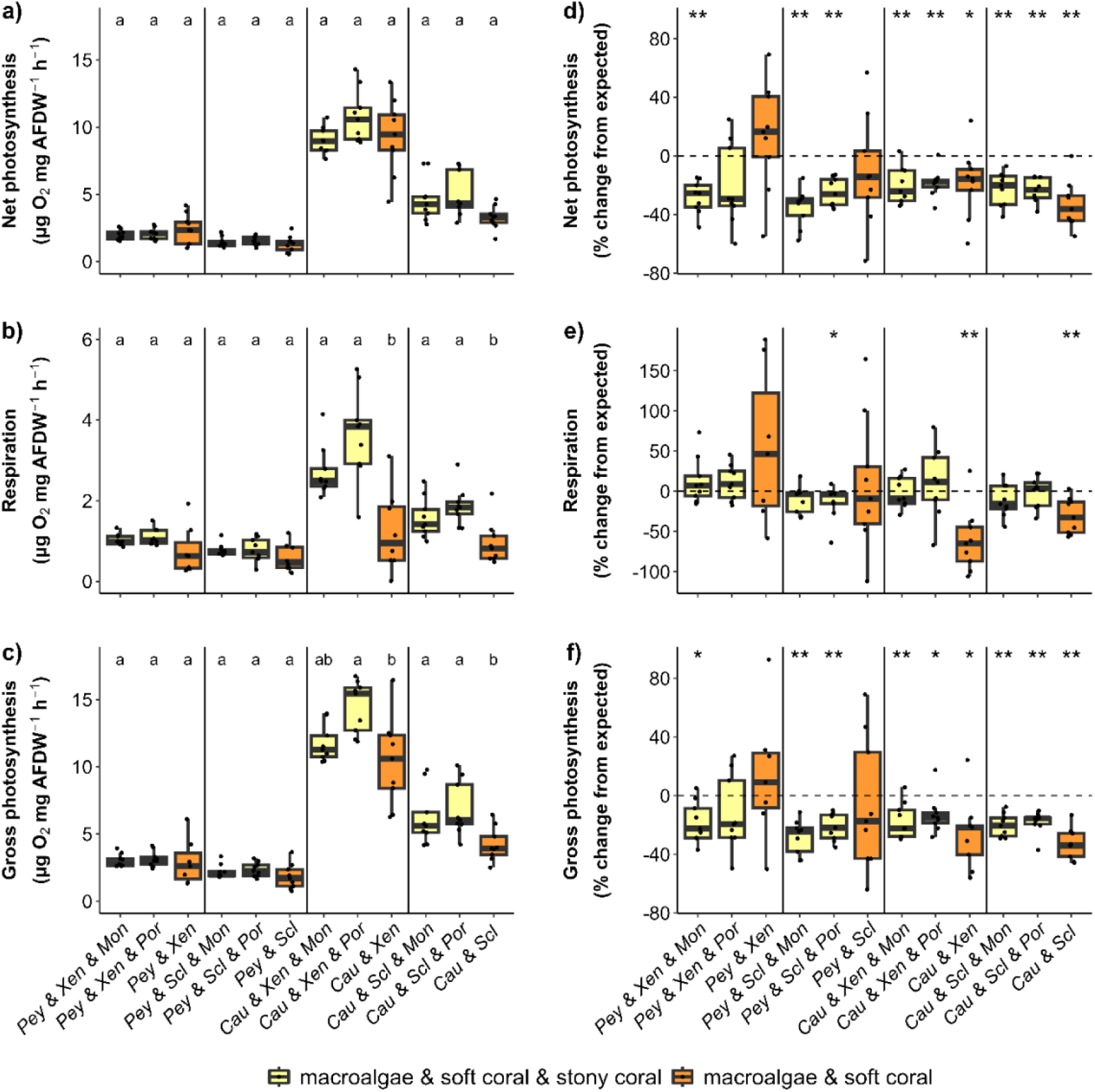
The absence of stony corals affects community productivity. This is shown as net photosynthesis (a), respiration (b), and gross photosynthesis (c) in absolute values and as change from expected (in %) of net photosynthesis (d), respiration (e), and gross photosynthesis (f). Species abbreviations: Mon = *Montipora*, Por = *Porites*, Xen = *Xenia*, Scl = *Sclerophytum*, Cau = *Caulerpa,* Pey = *Peyssonnelia*. Boxes indicate the first and third quartiles, and whiskers indicate ± 1.5 IQR. Letters indicate significant differences between soft coral & macroalgae incubations of the same species, with and without stony corals (a-c; p < 0.05). Asterisks indicate significant difference from expected (e-h; * p < 0.05, ** p < 0.01, *** p < 0.001; Tab. S6 & Tab. S7).

In relation to the expected community productivity based on monocultures of the individual species of that community, the combination of all three organism groups generally induced a reduction in photosynthesis, while respiration remained unaffected (Fig. 5 d-f, Tab. S7). Interestingly, the absence of the stony corals led to significantly lower photosynthesis and respiration than expected in the incubations with *C. brachypus*, with the strongest reduction in gross photosynthesis occurring with *S. wanannensis* (33 % ± 11 SD). In contrast, incubations with *Peyssonnelia* sp. were in a range with expected values, and interactions between *Peyssonnelia* sp. and *X. umbellata* even triggered a positive productivity trend.

### Experiment 2

#### Effect of water-mediated interactions on photosynthetic efficiency across species

To infer individual contributions to changes in community productivity, the photosynthetic efficiency of each organism was compared between monoculture and polyculture (combinations of two or three species, Fig. 6). Overall, photosynthetic efficiency was largely unaffected, with significant decreases or increases observed in only a few cases and always small in magnitude. Photosynthetic efficiency of both stony corals tended to decline in the presence of other organisms, with a single significant reduction (p = 0.01) observed in *M. digitata* incubated with *X. umbellata* (Fig. 6a, Tab. S8). Interestingly this negative effect was not observed when a third organism group was added. *P. rus* exhibited a negative trend with both soft corals, yet not with the two macroalgae or the triple mix (Fig. 6b). Photosynthetic efficiency of both soft corals remained unaffected by neighbouring organisms (Fig. 6c-d, Tab. S8). Macroalgae exhibited the strongest effects in photosynthetic efficiency when combined with other organisms. *Peyssonnelia* sp. significantly upregulated (p < 0.01) its photosynthetic efficiency in all four combinations with one other organism, with the highest increase in combination with *P. rus*. The same positive trend was observed in combinations with two other organisms (Fig. 6e). In contrast, *C. brachypus* significantly downregulated (p = 0.03) its photosynthetic efficiency in the presence of *M. digitata* and showed a negative trend with the two soft corals (Fig. 6f, Tab. S8). Yet, a positive trend was observed during the incubations with *P. rus* and the triple mixes.

**Figure 6.**
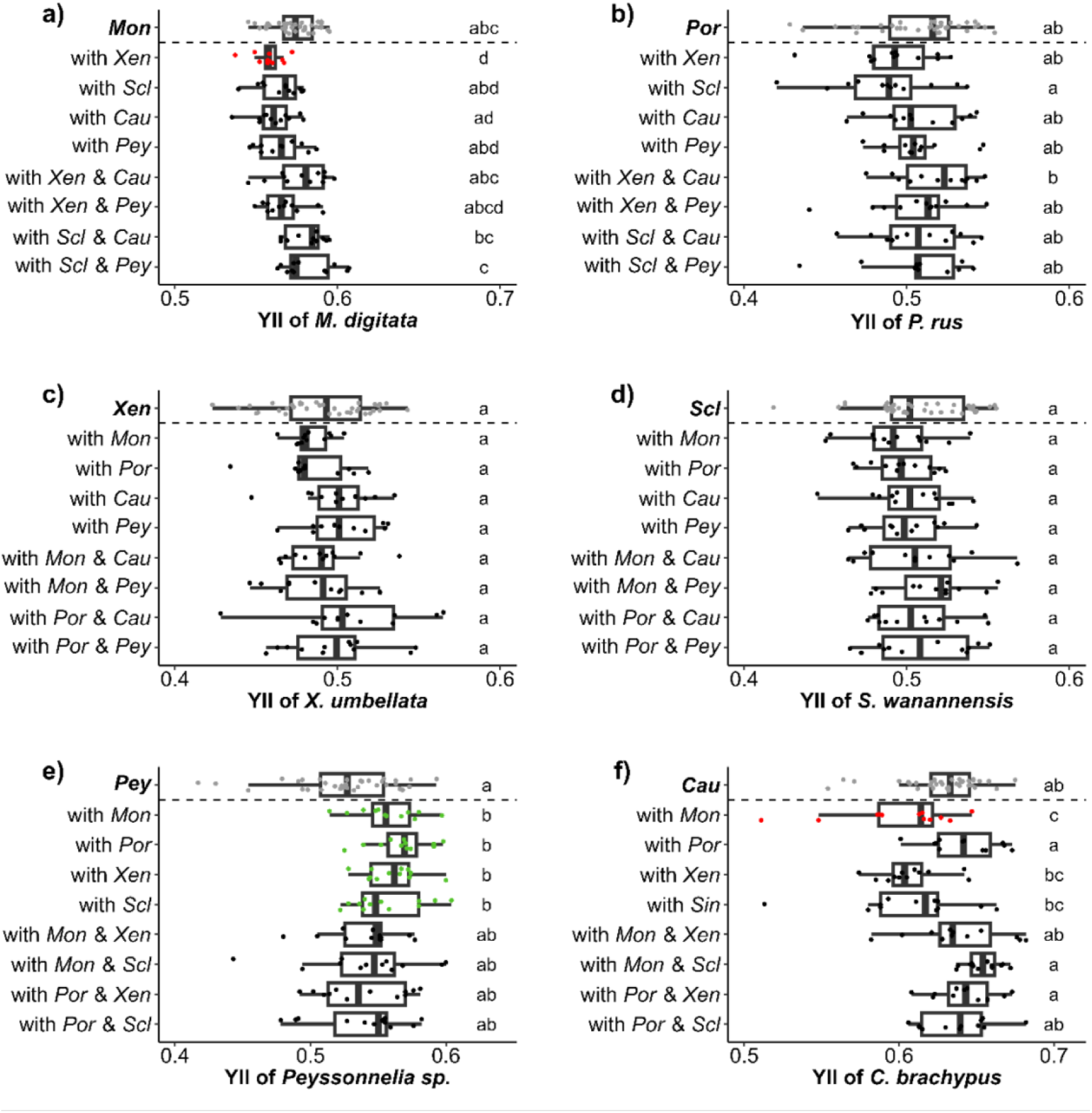
Influence of benthic reef neighbours on photosynthetic efficiency (YII) per species. Shown for *M. digitata* (Mon; a), *P. rus* (Por; b), *X. umbellata* (Xen; c), *S. wanannensis* (Scl; d), *Peyssonnelia* sp. (Pey; e), and *C. brachypus* (Cau; f). Boxes indicate the first and third quartiles, and whiskers indicate ± 1.5 IQR. Letters indicate significant differences between the various incubations (p < 0.05). Red datapoints indicate the polyculture incubations in which species show a significantly lower, green a significantly higher photosynthetic efficiency compared to their monocultures, which are shown in light grey.

## Discussion

Our controlled incubation experiments reveal that the negative feedback loop during shifts from stony coral to soft coral or macroalgae-dominated reefs may be partly driven by water-mediated interactions among the organisms. We observed a general decline in overall community productivity concurrent with reduced stony coral biomass. Specifically, stony corals exhibited decreased productivity in the presence of soft corals, while soft corals maintained productivity. Moreover, one macroalgae (*Peyssonnelia sp.*) increased productivity in the presence of other organisms, suggesting a competitive advantage. These patterns may partially explain the accelerated loss of stony corals once a critical biomass threshold is surpassed, contributing to the negative feedback mechanism driving the rapid degradation of coral reefs during such phase shifts.

### Baseline productivity varies between species

Species differed markedly in productivity, providing the baseline against which interaction effects in mixed communities can be interpreted. *C. brachypus* was most productive across all parameters, consistent with previous studies on *Caulerpa* spp. (Collado-Vides & Robledo, 1999; Terada et al., 2021). Following *C. brachypus*, the stony corals showed the highest productivity, followed by the two soft corals, and the least productive macroalga *Peyssonnelia* sp. (Fig. 3). The productivity patterns for the stony corals (Vetter et al., 2025) and soft corals (Fabricius & Klumpp, 1995) were also consistent with prior studies. Notably, *Peyssonnelia* sp. deviated from expected ranges, displaying low photosynthesis and calcification, probably due to its complex morphology with overarching plates (Pueschel & Saunders, 2009), complicating normalization. Variations in fragment thickness likely increased ash-free dry weight (AFDW) without enhancing the light absorbing surface area. This assumption is supported by the photosynthetic efficiency measurements, which placed it third highest among the six species, aligning with previous findings (Yıldız, 2018). Yet, these challenges did not affect the comparison between mono- and polyculture incubations, as the same *Peyssonnelia* fragments were consistently compared with each other.

### Phase shifts affect productivity

We show that community productivity is strongly governed by biomass ratios, with relative declines in productivity as soft coral or macroalgal biomass increases, exceeding the reduction anticipated from monoculture performance alone (Fig. 4). Declines in productivity with rising soft coral or macroalgal biomass suggest inhibitory interactions affecting at least one species of the community, likely mediated via allelopathic or microbial processes (Briggs et al., 2021; Clements & Hay, 2023; Haas et al., 2013). Our findings are in line with previous water-mediated interaction studies, that have shown that *Caulerpa taxifolia* had a negative effect on the stony corals *Turbinaria peltata* (Fu et al., 2023) and *Acropora hyacinthus* (Fu et al., 2022), while soft corals have been reported to reduce stony coral photosynthesis (Engelhardt et al., 2023; Smith et al., 2006). *Peyssonnelia* sp. exerted weaker impacts, which may be explained by a lack of allelopathic compounds in this species (Kase et al., 2020; Maida et al., 2001). To date, very few studies have addressed the possibility, that the observed changes in community productivity may also be driven by shifts in the productivity of the soft corals or macroalgae themselves. However, Andrade Rodríguez (2018) demonstrated that not only was gene expression in the stony coral *Porites cylindrica* affected by water-mediated interactions with other species, but gene expression was also induced in the soft coral *Lobophytum pauciflorum*, suggesting bidirectional physiological responses.

While interaction effects were significant, their absolute magnitude remained moderate. For example, incubations containing *C. brachypus* maintained some of the highest absolute productivity values, even though relative productivity compared to monoculture expectations declined. This is consistent with previous studies, which have documented water-mediated interactions affecting productivity, but suggest that their overall importance under natural environmental conditions is relatively low (Chadwick & Morrow, 2011; Clements & Hay, 2023; Fu et al., 2023; Sammarco et al., 1983). Nevertheless, communities dominated by stony corals deviated least from monoculture expectations, highlighting the disproportionate metabolic destabilization that may occur as stony coral dominance declines, potentially triggering cascading effects.

### Loss of stony corals reshapes benthic interactions

Community productivity changes may be governed more by the presence of stony corals than by which species are present. When all three functional groups (stony corals, soft corals, and macroalgae) were incubated together, community productivity consistently was reduced compared to monoculture expectations (Fig. 5). Assemblages containing the same two soft corals and macroalgae but different stony corals (*M. digitata* or *P. rus*) showed comparable declines, suggesting that species identity mattered less than stony coral presence itself. When stony corals were absent, deviations from expected often became more pronounced, underscoring the need to consider interactions among non-scleractinian taxa as well. In *C. brachypus*–soft coral combinations, both absolute and expected productivity declined significantly, suggesting that *C. brachypus* may employ a different competitive strategy in the presence of only soft coral neighbours, shifting between chemical defence and growth (Pergent et al., 2008). Conversely, incubations with *Peyssonnellia* sp. exhibited reduced productivity when combined with both soft and stony corals yet showed no deviation from expected productivity when stony corals were absent, suggesting that stony corals may be a strong competitor for them. These results indicate that the presence of stony corals may modulate the direction and strength of interactions among other benthic organisms.

The short-term productivity responses we observed align with broader evidence of long-term shifts in community productivity following phase shifts to soft corals and macroalgae (Done, 1992; McManus & Polsenberg, 2004; Mumby, 2009; Norström et al., 2009). Such changes reflect a complex interplay among host organisms, their associated microbiomes, released metabolites, and nutrient cycling (Norström et al., 2009). Our findings suggest that even at the microscale, within small community patches, species constantly adjust their productivity in response to changes of their neighbours.

### Productivity changes reveal nuanced interspecific effects in reef communities

At the species level, photosynthetic efficiency patterns reveal subtle but informative trends, from which several conclusions can be drawn. First, soft coral photosynthetic efficiency was not affected, which hints that the downregulation in photosynthesis of the first experiment was due to stony corals (Fig. 4) and macroalgae (Fig. 5). This is consistent with evidence that soft coral exometabolomes are highly responsive to neighbours (Weber et al., 2022; Engelhardt, 2025), potentially releasing allelopathic or signaling compounds that influence microbial communities and thereby indirectly reduce overall productivity. The lack of significant changes in their own photosynthetic efficiency may indicate that soft corals tolerate these interactions physiologically, while impacting the productivity of more sensitive neighbours. Second, both stony corals showed small negative responses to neighbours. This pattern aligns with our previous finding that *P. rus* and *M. digitata* tended to respond negatively to neighbours, whereas other stony coral species sometimes showed positive effects (Vetter et al., 2025), emphasizing the importance of testing broader species combinations. In our study, as in Vetter et al. (2025), *P. rus* reacted less strongly than *M. digitata* which may reflect its stress-tolerant nature, consistent with known resilience to temperature and nutrient disturbances (Donner & Carilli, 2019). Finally, *Peyssonnelia* sp. consistently increased photosynthetic efficiency when grown with other species, which may explain some of the non-significant net changes in photosynthesis in mixed incubations (Figs. 4 & 5), as productivity declines in other taxa were partly offset by gains in *Peyssonnelia* sp. Given that *Peyssonnelia* sp. is recognized as a competitive and stress-tolerant genus capable of overgrowing reefs (Edmunds et al., 2023; Yıldız, 2018), these results suggest a potential mechanism contributing to its ecological success. Field studies typically report reduced coral photosynthetic efficiency only under direct contact with macroalgae (Clements & Hay, 2023; de Nys et al., 1991; Rasher & Hay, 2010). In contrast, our results demonstrate that water-mediated interactions alone elicit measurable shifts, which were particularly pronounced in *Peyssonnelia* sp. These water-mediated effects should therefore receive more attention, even if strong currents and rapid dilution in the field will likely dampen their ecological impact. Given that elevated macroalgal cover is associated with increased microbial loads and disease prevalence (Haas et al., 2016), productivity shifts may signal early, sublethal stress responses, potentially diverting energy away from immunity and heightening vulnerability to disturbances.

### Roles of microbiomes and metabolites in shaping productivity

Integrating our findings with previous work on microbial and chemical interactions provides mechanistic insights. Using the same species, Wiederkehr et al. (2025) found that a less diverse community reshapes microbial networks. Stony coral microbiomes become less connected, whereas those of soft corals, macroalgae become more tightly linked (Wiederkehr et al., 2025). These changes are not merely a reduction in microbial diversity but a restructuring of microbial interactions that regulate ecosystem function. Thus biodiversity particularly influences the acquisition of novel microbes, a process that may underlie holobiont adaptability to environmental stressors (Voolstra & Ziegler, 2020) depending on the microbiome flexibility of the species (Ziegler et al., 2019).

These microbial dynamics are closely intertwined with the reef exometabolome, as the composition and release of benthic metabolites depend not only on species identity but also on the surrounding community context (Engelhardt et al. 2025; Weber et al., 2022). Soft corals and macroalgae release more metabolites compared with stony corals, and specific exometabolites are only produced in mixed assemblages, highlighting active biochemical responses to neighboring taxa (Weber et al., 2022; Engelhardt et al., 2025). These metabolites may then affect photosynthesis in specific hosts (Weber et al., 2022).

### Future directions

Together, our findings show that species within small community patches continuously adjust their productivity in response to changes in the presence and prevalence of their neighbours. Even subtle changes in community composition may disproportionately alter productivity dynamics, highlighting their potential role in mediating coral reef phase shifts. The water column serves as a central conduit for these interactions, yet its inherent variability makes them challenging to study (Shakya & Allgeier, 2023). Future research should aim to disentangle the mechanisms underlying these effects, clarifying which productivity changes are driven by the host organisms and their associated microbial communities, and how these interactions are influenced by anthropogenic stressors. For example, ocean acidification increases productivity of macroalgae incl. *Peyssonnelia* sp. (Yıldız, 2018), while it typically decreases that of stony corals (Anthony et al., 2008), and it is important to examine these responses in the context of interactions with neighbouring species. Similarly, *Xenia* spp. and other soft corals typically benefit from high tolerance to temperature and nutrient stress (Mezger et al., 2022) compared to stony corals, and feedback loops under these scenarios remain to be investigated. Thus, experiments under climate change scenarios are particularly needed to predict how interactions may shift, and their outcomes should be integrated into reef models and restoration planning to anticipate the effects of climate change and pollution.

## Statements and Declarations

### Author contributions

J.V., K.E.E., and M.Z. conceived the study and designed the experiments. J.V., K.E.E., A.D., and F.W-Z. collected data. J.V., K.E.E. and A.D. analysed and curated data. J.V. wrote the manuscript with contributions from M.Z. and K.E.E. All authors read and approved the final manuscript.

### Data and Code availability

The datasets generated and analyzed during this study as well as the R scripts are available in GitHub https://github.com/JanaVetter/PhaseShiftExp_manuscript

## References

Aceret, T. L., Sammarco, P. W., & Coll, J. C. (1995). Toxic effects of alcyonacean diterpenes on scleractinian corals. Journal of Experimental Marine Biology and Ecology, 188(1), 63–78. 10.1016/0022-0981(94)00186-H

Andrade Rodríguez, N. A. (2018). Non-contact competition between soft and hard corals: A transcriptomic perspective. 10.25903/5BDA8F54CF401

Anthony, K. R. N., Kline, D. I., Diaz-Pulido, G., Dove, S., & Hoegh-Guldberg, O. (2008). Ocean acidification causes bleaching and productivity loss in coral reef builders. Proceedings of the National Academy of Sciences, 105(45), 17442–17446. 10.1073/pnas.0804478105

Anton, A., Randle, J. L., Garcia, F. C., Rossbach, S., Ellis, J. I., Weinzierl, M., & Duarte, C. M. (2020). Differential thermal tolerance between algae and corals may trigger the proliferation of algae in coral reefs. Global Change Biology, 26(8), 4316–4327. 10.1111/gcb.15141

Briggs, A. A., Brown, A. L., & Osenberg, C. W. (2021). Local versus site-level effects of algae on coral microbial communities. Royal Society Open Science, 8(9), 210035. 10.1098/rsos.210035

Chadwick, N. E., & Morrow, K. M. (2011). Competition Among Sessile Organisms on Coral Reefs. In Z. Dubinsky & N. Stambler (Eds.), Coral Reefs: An Ecosystem in Transition (pp. 347–371). Springer Netherlands. 10.1007/978-94-007-0114-4_20

Chisholm, J. R. M., & Gattuso, J.-P. (1991). Validation of the alkalinity anomaly technique for investigating calcification of photosynthesis in coral reef communities. Limnology and Oceanography, 36(6), 1232–1239. 10.4319/lo.1991.36.6.1232

Clements, C. S., & Hay, M. E. (2023). Disentangling the impacts of macroalgae on corals via effects on their microbiomes. Frontiers in Ecology and Evolution, 11. https://www.frontiersin.org/articles/10.3389/fevo.2023.1083341

Collado-Vides, L., & Robledo, D. (1999). MORPHOLOGY AND PHOTOSYNTHESIS OF *CAULERPA* (CHLOROPHYTA) IN RELATION TO GROWTH FORM. Journal of Phycology, 35(2), 325–330. 10.1046/j.1529-8817.1999.3520325.x

Done, T.J. (1992). Phase shifts in coral reef communities and their ecological significance. Hydrobiologia, 247, 121–132.

Dumay, O., Fernandez, C., & Pergent, G. (2002). Primary production and vegetative cycle in *Posidonia oceanica* when in competition with the green algae *Caulerpa taxifolia* and *Caulerpa racemosa*. Journal of the Marine Biological Association of the United Kingdom, 82(3), 379–387. 10.1017/S0025315402005611

Edmunds, P. J., Schils, T., & Wilson, B. (2023). The rising threat of peyssonnelioid algal crusts on coral reefs. Current Biology, 33(21), R1140–R1141. 10.1016/j.cub.2023.08.097

Engelhardt, K. E., Vetter, J., Wöhrmann-Zipf, F., Dietzmann, A., Proll, F. M., Reifert, H., Schüll, I., Stahlmann, M., & Ziegler, M. (2023). Contact-free impacts of sessile reef organisms on stony coral productivity. Communications Earth & Environment, 4(1), Article 1. 10.1038/s43247-023-01052-5

Fabricius, K. E., & Klumpp, D. W. (1995). Widespread mixotrophy in reef-inhabiting soft corals: The influence of depth, and colony expansion and contraction on photosynthesis. Marine Ecology Progress Series, 125, 195–204. 10.3354/meps125195

Fong, J., Deignan, L. K., Bauman, A. G., Steinberg, P. D., McDougald, D., & Todd, P. A. (2020). Contact- and Water-Mediated Effects of Macroalgae on the Physiology and Microbiome of Three Indo-Pacific Coral Species. Frontiers in Marine Science, 6. 10.3389/fmars.2019.00831

Fu, J., Zhou, J., Zhou, J., Zhang, Y., & Liu, L. (2023). Competitive effects of the macroalga Caulerpa taxifolia on key physiological processes in the scleractinian coral Turbinaria peltata under thermal stress. PeerJ, 11, e16646. 10.7717/peerj.16646

Fu, Zhou, J., Zhang, Y.-P., & Liu, L. (2022). Effects of Caulerpa taxifolia on Physiological Processes and Gene Expression of Acropora hyacinthus during Thermal Stress. Biology, 11(12), Article 12. 10.3390/biology11121792

Graham, N. A. J., Jennings, S., MacNeil, M. A., Mouillot, D., & Wilson, S. K. (2015). Predicting climate-driven regime shifts versus rebound potential in coral reefs. Nature, 518, 94–97.

Haas, A. F., Fairoz, M. F. M., Kelly, L. W., Nelson, C. E., Dinsdale, E. A., Edwards, R. A., Giles, S., Hatay, M., Hisakawa, N., Knowles, B., Lim, Y. W., Maughan, H., Pantos, O., Roach, T. N. F., Sanchez, S. E., Silveira, C. B., Sandin, S., Smith, J. E., & Rohwer, F. (2016). Global microbialization of coral reefs. Nature Microbiology, 1(6), 16042. 10.1038/nmicrobiol.2016.42

Haas, A. F., Nelson, C. E., Rohwer, F., Wegley-Kelly, L., Quistad, S. D., Carlson, C. A., Leichter, J. J., Hatay, M., & Smith, J. E. (2013). Influence of coral and algal exudates on microbially mediated reef metabolism. PeerJ, 1, e108. 10.7717/peerj.108

Hothorn, T., Bretz, F., & Westfall, P. (2008). Simultaneous Inference in General Parametric Models. Biometrical Journal, 50(3), 346–363.

Huey, R. B., Carlson, M., Crozier, L., Frazier, M., Hamilton, H., Harley, C., Hoang, A., & Kingsolver, J. G. (2002). Plants Versus Animals: Do They Deal with Stress in Different Ways?1. Integrative and Comparative Biology, 42(3), 415–423. 10.1093/icb/42.3.415

Kase, A. G. O., Calumpong, H., & Rupidara, A. (2020). Secondary metabolites of some varieties of Caulerpa species. IOP Conference Series: Materials Science and Engineering, 823(1), 012041. 10.1088/1757-899X/823/1/012041

Maida, M., Sammarco, P. W., & Coll, J. C. (2001). Effects of Soft Corals on Scleractinian Coral Recruitment. II: Allelopathy, Spat Survivorship and Reef Community Structure. Marine Ecology, 22(4), 397–414. 10.1046/j.1439-0485.2001.01709.x

McManus, J. W., & Polsenberg, J. F. (2004). Coral–algal phase shifts on coral reefs: Ecological and environmental aspects. Progress in Oceanography, 60(2–4), 263–279. 10.1016/j.pocean.2004.02.014

Mezger, S. D., Klinke, A., Tilstra, A., El-Khaled, Y. C., Thobor, B., & Wild, C. (2022). The widely distributed soft coral Xenia umbellata exhibits high resistance against phosphate enrichment and temperature increase. Scientific Reports, 12(1), 22135. 10.1038/s41598-022-26325-5

Mumby, P. J. (2009). Phase shifts and the stability of macroalgal communities on Caribbean coral reefs. Coral Reefs, 28(3), 761–773. 10.1007/s00338-009-0506-8

Norström, A., Nyström, M., Lokrantz, J., & Folke, C. (2009). Alternative states on coral reefs: Beyond coral–macroalgal phase shifts. Marine Ecology Progress Series, 376, 295–306. 10.3354/meps07815

Pergent, G., Boudouresque, C.-F., Dumay, O., Pergent-Martini, C., & Wyllie-Echeverria, S. (2008). Competition between the invasive macrophyte Caulerpa taxifolia and the seagrass Posidonia oceanica: Contrasting strategies. BMC Ecology, 8(1), 20. 10.1186/1472-6785-8-20

Pinheiro, J., Bates, D., DebRoy, S., Sarkar, D., & R Core Team. (2020). *_**nlme: Linear and Nonlinear Mixed Effects Models_. R package version 3.1-149* [Computer software]. R package version 3.1-149. https://CRAN.R-project.org/package=nlme

Pueschel, C. M., & Saunders, G. W. (2009). *Ramicrusta textilis* sp. Nov. (Peyssonneliaceae, Rhodophyta), an anatomically complex Caribbean alga that overgrows corals. Phycologia, 48(6), 480–491. 10.2216/09-04.1

Pupier, C. A., Bednarz, V. N., & Ferrier-Pagès, C. (2018). Studies With Soft Corals – Recommendations on Sample Processing and Normalization Metrics. Frontiers in Marine Science, 5. 10.3389/fmars.2018.00348

R Core Team. (2023). R: A language and environment for statistical computing. R Foundation for Statistical Computing [Computer software]. https://www.R-project.org/

Rades, M., Schubert, P., Ziegler, M., Kröckel, M., & Reichert, J. (2022). Building plan for a temperature-controlled multi-point stirring incubator. Protocols.Io 10. 10.17504/protocols.io.dm6gpb34dlzp/v1

Reaka-Kudla, M. L. (1997). The Global Biodiversity of Coral Reefs: A Comparison with Rain Forests. In Biodiversity II: Understanding and protecting our biological resources. (pp. 83–108). The National Academie of Science.

Riegl, B., & Piller, W. E. (1999). Coral frameworks revisited–reefs and coral carpets in the northern Red Sea. Coral Reefs, 18(3), 241–253. 10.1007/s003380050188

Rodriguez, N. A., Moya, A., Jones, R., Miller, D. J., & Cooke, I. (2021). The Significance of Genotypic Diversity in Coral Competitive Interaction: A Transcriptomic Perspective. Frontiers in Ecology and Evolution. 10.26181/60ed15a645426

Roff, G., Chollett, I., Doropoulos, C., Golbuu, Y., Steneck, R. S., Isechal, A. L., van Woesik, R., & Mumby, P. J. (2015). Exposure-driven macroalgal phase shift following catastrophic disturbance on coral reefs. Coral Reefs, 34(3), 715–725. 10.1007/s00338-015-1305-z

Roth, F., RAdecker, N., Carvalho, S., Duarte, C. M., Saderne, V., Anton, A., Silva, L., Calleja, M. Ll., MorÁn, X. A. G., Voolstra, C. R., Kürten, B., Jones, B. H., & Wild, C. (2021). High summer temperatures amplify functional differences between coral- and algae-dominated reef communities. Ecology, 102(2), e03226. 10.1002/ecy.3226

Sammarco, P. W., Coll, J. C., La Barre, S., & Willis, B. (1983). Competitive strategies of soft corals (Coelenterata: Octocorallia): Allelopathic effects on selected scleractinian corals. Coral Reefs, 1(3), 173–178. 10.1007/BF00571194

Schneider, K., & Erez, J. (2006). The effect of carbonate chemistry on calcification and photosynthesis in the hermatypic coral Acropora eurystoma. Limnology and Oceanography, 51(3), 1284–1293. 10.4319/lo.2006.51.3.1284

Shakya, A. W., & Allgeier, J. E. (2023). Water column contributions to coral reef productivity: Overcoming challenges of context dependence. Biological Reviews, 98(5), 1812–1828. 10.1111/brv.12984

Smith, J. E., Shaw, M., Edwards, R. A., Obura, D., Pantos, O., Sala, E., Sandin, S. A., Smriga, S., Hatay, M., & Rohwer, F. L. (2006). Indirect effects of algae on coral: Algae-mediated, microbe-induced coral mortality. Ecology Letters, 9(7), 835–845. 10.1111/j.1461-0248.2006.00937.x

Terada, R., Takaesu, M., Borlongan, I. A., & Nishihara, G. N. (2021). The photosynthetic performance of a cultivated Japanese green alga Caulerpa lentillifera in response to three different stressors, temperature, irradiance, and desiccation. Journal of Applied Phycology, 33(4), 2547–2559. 10.1007/s10811-021-02439-7

Thurber, R. V., Burkepile, D. E., Correa, A. M. S., Thurber, A. R., Shantz, A. A., Welsh, R., Pritchard, C., & Rosales, S. (2012). Macroalgae Decrease Growth and Alter Microbial Community Structure of the Reef-Building Coral, Porites astreoides. PLOS ONE, 7(9), e44246. 10.1371/journal.pone.0044246

Vetter, J., Reichert, J., Dietzmann, A., Hahn, L., Lang, A. E., Puntin, G., & Ziegler, M. (2025). Species identity and composition affect the productivity of stony corals. Coral Reefs. 10.1007/s00338-025-02748-0

Voolstra, C. R., & Ziegler, M. (2020). Adapting with Microbial Help: Microbiome Flexibility Facilitates Rapid Responses to Environmental Change. *BioEssays: News and Reviews in Molecular*, Cellular and Developmental Biology, 42(7), e2000004. 10.1002/bies.202000004

Wickham, H., Averick, M., Bryan, J., Chang, W., McGowan, L. D., & François, R. (2019). *“*Welcome to the tidyverse*.”* [Computer software]. J. Open Source Softw., 4(43), 1686.

Wiederkehr, F., Engelhardt, K. E., Vetter, J., Ruscheweyh, H.-J., Salazar, G., O’Brien, J., Priest, T., Ziegler, M., & Sunagawa, S. (2025). Host-level biodiversity shapes the dynamics and networks within the coral reef microbiome. ISME Communications, 5(1), ycaf097. 10.1093/ismeco/ycaf097

Yıldız, G. (2018). Physiological Responses of the Mediterranean Subtidal Alga Peyssonnelia squamaria to Elevated CO2. Ocean Science Journal, 53(4), 691–698. 10.1007/s12601-018-0044-9

Ziegler, M., Grupstra, C. G. B., Barreto, M. M., Eaton, M., BaOmar, J., Zubier, K., Al-Sofyani, A., Turki, A. J., Ormond, R., & Voolstra, C. R. (2019). Coral bacterial community structure responds to environmental change in a host-specific manner. Nature Communications, 10(1), 3092. 10.1038/s41467-019-10969-5

