## Supplemental Material for "Water-mediated productivity dynamics in shifting coral reef communities"

### Supplementary Tables

**Species abbreviations in supplementary tables: *Montipora digitata* (Mon), *Porites rus* (Por), *Xenia umbellata* (Xen), *Sclerophytum wanannensis* (Scl), *Caulerpa brachypus* (Cau), *Peyssonnelia* sp. (Pey)**

**Tab. S1 CITES permit numbers for the used stony coral colonies**

| Family | Species | CITES # | Origin |
|---|---|---|---|
| Poritidae | *P. rus* | 15-SA-000885-PD | Red Sea |
| Acroporidae | *M. digitata* | FSHQ/340/14 | Indopacific |

**Tab. S2 One way ANOVA of the monoculture assemblages, including coral fragment ID as random factor, comparing the mean productivity of the six organisms**, followed by a Tukey post hoc test with Bonferroni correction (Fig. 3). Significant changes ($p < 0.05$) are indicated in bold. Net photosynthesis and gross photosynthesis were log-transformed. Three photosynthetic values, six calcification values and two PAM values were found to be outliers and removed from the dataset.

| | One-way Anova | | | | | Tukey with Bonferroni correction | | | | |
|---|---|---|---|---|---|---|---|---|---|---|
| Variable | | #DF | den DF | F value | p-value | Groups | estimate | Std. error | Z value | p adj. |
| net photo-synthesis | Int. | 1 | 107 | 2898 | **<.001** | Por - Mon | 0.38 | 0.12 | 3.1 | **0.026** |
| | Species | 5 | 66 | 193 | **<.001** | Xen - Mon | -0.19 | 0.12 | -1.6 | 0.602 |
| | | | | | | Scl - Mon | -1.18 | 0.12 | -9.6 | **<.001** |
| | | | | | | Pey - Mon | -1.66 | 0.12 | -13.5 | **<.001** |
| | | | | | | Cau - Mon | 1.20 | 0.11 | 11.3 | **<.001** |
| | | | | | | Xen - Por | -0.57 | 0.12 | -4.7 | **<.001** |
| | | | | | | Scl - Por | -1.56 | 0.12 | -12.7 | **<.001** |
| | | | | | | Pey - Por | -2.04 | 0.12 | -16.5 | **<.001** |
| | | | | | | Cau - Por | 0.82 | 0.11 | 7.8 | **<.001** |
| | | | | | | Scl - Xen | -0.98 | 0.12 | -8.0 | **<.001** |
| | | | | | | Pey - Xen | -1.47 | 0.12 | -11.9 | **<.001** |
| | | | | | | Cau - Xen | 1.39 | 0.11 | 13.2 | **<.001** |
| | | | | | | Pey - Scl | -0.48 | 0.12 | -3.9 | **0.001** |
| | | | | | | Cau - Scl | 2.38 | 0.11 | 22.5 | **<.001** |
| | | | | | | Cau - Pey | 2.86 | 0.11 | 26.9 | **<.001** |
| res-piration | Int. | 1 | 105 | 536 | **<.001** | Por - Mon | 1.56 | 0.38 | 4.1 | **<.001** |
| | Species | 5 | 66 | 37 | **<.001** | Xen - Mon | -0.50 | 0.38 | -1.3 | 0.775 |
| | | | | | | Scl - Mon | -1.48 | 0.38 | -3.9 | **0.001** |
| | | | | | | Pey - Mon | -2.10 | 0.39 | -5.4 | **<.001** |
| | | | | | | Cau - Mon | 1.64 | 0.34 | 4.9 | **<.001** |
| | | | | | | Xen - Por | -2.06 | 0.38 | -5.4 | **<.001** |
| | | | | | | Scl - Por | -3.04 | 0.38 | -8.1 | **<.001** |
| | | | | | | Pey - Por | -3.66 | 0.39 | -9.4 | **<.001** |
| | | | | | | Cau - Por | 0.08 | 0.34 | 0.2 | 1.000 |
| | | | | | | Scl - Xen | -0.99 | 0.38 | -2.6 | 0.092 |

| | One-way Anova | | | | | Tukey with Bonferroni correction | | | | |
|---|---|---|---|---|---|---|---|---|---|---|
| Variable | | #DF | den DF | F value | p-value | Groups | estimate | Std. error | Z value | p adj. |
| | | | | | | Pey - Xen | -1.61 | 0.39 | -4.1 | **0.001** |
| | | | | | | Cau - Xen | 2.14 | 0.34 | 6.3 | **<.001** |
| | | | | | | Pey - Scl | -0.62 | 0.39 | -1.6 | 0.603 |
| | | | | | | Cau - Scl | 3.12 | 0.34 | 9.3 | **<.001** |
| | | | | | | Cau - Pey | 3.74 | 0.35 | 10.7 | **<.001** |
| gross photo-synthesis | Int. | 1 | 107 | 4283 | **<.001** | Por - Mon | 0.44 | 0.12 | 3.7 | **0.002** |
| | Species | 5 | 66 | 208 | **<.001** | Xen - Mon | -0.22 | 0.12 | -1.9 | 0.398 |
| | | | | | | Scl - Mon | -1.11 | 0.12 | -9.6 | **<.001** |
| | | | | | | Pey - Mon | -1.83 | 0.12 | -15.7 | **<.001** |
| | | | | | | Cau - Mon | 1.05 | 0.10 | 10.5 | **<.001** |
| | | | | | | Xen - Por | -0.66 | 0.12 | -5.7 | **<.001** |
| | | | | | | Scl - Por | -1.55 | 0.12 | -13.3 | **<.001** |
| | | | | | | Pey - Por | -2.26 | 0.12 | -19.4 | **<.001** |
| | | | | | | Cau - Por | 0.62 | 0.10 | 6.2 | **<.001** |
| | | | | | | Scl - Xen | -0.89 | 0.12 | -7.7 | **<.001** |
| | | | | | | Pey - Xen | -1.61 | 0.12 | -13.8 | **<.001** |
| | | | | | | Cau - Xen | 1.27 | 0.10 | 12.7 | **<.001** |
| | | | | | | Pey - Scl | -0.71 | 0.12 | -6.1 | **<.001** |
| | | | | | | Cau - Scl | 2.17 | 0.10 | 21.7 | **<.001** |
| | | | | | | Cau - Pey | 2.88 | 0.10 | 28.6 | **<.001** |
| calci-fication | Int. | 1 | 101 | 129 | **<.001** | Por - Mon | 0.01 | 0.01 | 0.9 | 0.941 |
| | Species | 5 | 65 | 45 | **<.001** | Xen - Mon | -0.11 | 0.01 | -8.1 | **<.001** |
| | | | | | | Scl - Mon | -0.11 | 0.01 | -7.9 | **<.001** |
| | | | | | | Pey - Mon | -0.12 | 0.01 | -8.8 | **<.001** |
| | | | | | | Cau - Mon | -0.12 | 0.01 | -9.6 | **<.001** |
| | | | | | | Xen - Por | -0.13 | 0.01 | -9.0 | **<.001** |
| | | | | | | Scl - Por | -0.12 | 0.01 | -8.8 | **<.001** |
| | | | | | | Pey - Por | -0.14 | 0.01 | -9.7 | **<.001** |

| | One-way Anova | | | | | Tukey with Bonferroni correction | | | | |
|---|---|---|---|---|---|---|---|---|---|---|
| Variable | | #DF | den DF | F value | p-value | Groups | estimate | Std. error | Z value | p adj. |
| | | | | | | Cau - Por | -0.13 | 0.01 | -10.8 | **<.001** |
| | | | | | | Scl - Xen | 0.00 | 0.01 | 0.2 | 1.000 |
| | | | | | | Pey - Xen | -0.01 | 0.01 | -0.6 | 0.992 |
| | | | | | | Cau - Xen | 0.00 | 0.01 | -0.1 | 1.000 |
| | | | | | | Pey - Scl | -0.01 | 0.01 | -0.8 | 0.963 |
| | | | | | | Cau - Scl | 0.00 | 0.01 | -0.4 | 0.999 |
| | | | | | | Cau - Pey | 0.01 | 0.01 | 0.6 | 0.991 |
| Y II | Int. | 1 | 142 | 45479 | **<.001** | Por - Mon | -0.07 | 0.01 | -7.8 | **<.001** |
| | Species | 5 | 66 | 73 | <.001 | Xen - Mon | -0.08 | 0.01 | -9.5 | **<.001** |
| | | | | | | Scl - Mon | -0.07 | 0.01 | -7.6 | **<.001** |
| | | | | | | Pey - Mon | -0.05 | 0.01 | -5.5 | **<.001** |
| | | | | | | Cau - Mon | 0.06 | 0.01 | 6.3 | **<.001** |
| | | | | | | Xen - Por | -0.01 | 0.01 | -1.7 | 0.558 |
| | | | | | | Scl - Por | 0.00 | 0.01 | 0.2 | 1.000 |
| | | | | | | Pey - Por | 0.02 | 0.01 | 2.2 | 0.243 |
| | | | | | | Cau - Por | 0.12 | 0.01 | 14.1 | **<.001** |
| | | | | | | Scl - Xen | 0.02 | 0.01 | 1.9 | 0.412 |
| | | | | | | Pey - Xen | 0.03 | 0.01 | 3.8 | **0.002** |
| | | | | | | Cau - Xen | 0.14 | 0.01 | 15.8 | **<.001** |
| | | | | | | Pey - Scl | 0.02 | 0.01 | 2.0 | 0.361 |
| | | | | | | Cau - Scl | 0.12 | 0.01 | 13.9 | **<.001** |
| | | | | | | Cau - Pey | 0.10 | 0.01 | 11.8 | **<.001** |

**Tab. S3 Productivity per species in monoculture**

| Group | Species | Productivity parameters (mean ± SD) | | | | |
|---|---|---|---|---|---|---|
| | | net photosynthesis ($\mu g\ O_2\ cm^{-2}\ h^{-1}$) | respiration ($\mu g\ O_2\ cm^{-2}\ h^{-1}$) | gross photosynthesis ($\mu g\ O_2\ cm^{-2}\ h^{-1}$) | calcification ($\mu mol\ CaCO_3\ cm^{-2}\ h^{-1}$) | Photosynthetic efficiency (YII) |
| Stony corals | *P. rus* | 88.8 ± 21.9 | 39.0 ± 13.9 | 127.8 ± 24.7 | 0.14 ± 0.06 | 0.506 ± 0.014 |
| | *M. digitata* | 61.6 ± 18.8 | 23.4 ± 10.3 | 85.1 ± 27.3 | 0.12 ± 0.05 | 0.574 ± 0.006 |
| Soft corals | *X. umbellata* | 51.6 ± 19.1 | 18.5 ± 12.9 | 7.0 ± 28.7 | 0.01 ± 0.02 | 0.491 ± 0.014 |
| | *S. wanannense* | 2.0 ± 0.9 | 0.9 ± 0.3 | 2.9 ± 10.8 | 0.01 ± 0.01 | 0.508 ± 0.015 |
| Macro algae | *Peyssonnelia* sp. | 1.3 ± 0.6 | 0.2 ± 0.2 | 14.2 ± 0.6 | 0.00 ± 0.02 | 0.525 ± 0.016 |
| | *C. brachypus* | 210.8 ± 80.8 | 39.3 ± 21.5 | 25.07 ± 98.4 | 0.01 ± 0.04 | 0.629 ± 0.012 |

**Tab. S4 Paired t-test with Bonferroni correction comparing measured community productivity between incubations with high stony coral biomass and those with higher soft coral or macroalgae biomass (Fig. 4 a-d).** Significant changes ($p < 0.05$) are indicated in bold

| Phase Shift incu-bations | Variable | Group1 | Group2 | n1 | n2 | statistic | df | p.adj |
|---|---|---|---|---|---|---|---|---|
| *Mon & Xen* | net photosynthesis | < stony corals | < soft corals | 9 | 9 | 1.440 | 8 | 0.188 |
| | respiration | < stony corals | < soft corals | 9 | 9 | 1.588 | 8 | 0.151 |
| | gross photosynthesis | < stony corals | < soft corals | 9 | 9 | 1.557 | 8 | 0.158 |
| | calcification | < stony corals | < soft corals | 9 | 9 | 4.671 | 8 | **0.002** |
| *Por & Xen* | net photosynthesis | < stony corals | < soft corals | 9 | 9 | 2.679 | 8 | **0.028** |
| | respiration | < stony corals | < soft corals | 9 | 9 | 1.354 | 8 | 0.213 |
| | gross photosynthesis | < stony corals | < soft corals | 9 | 9 | 3.000 | 8 | **0.017** |
| | calcification | < stony corals | < soft corals | 9 | 8 | 1.527 | 7 | 0.171 |
| *Mon & Scl* | net photosynthesis | < stony corals | < soft corals | 9 | 9 | 3.150 | 8 | **0.014** |
| | respiration | < stony corals | < soft corals | 9 | 9 | 0.686 | 8 | 0.512 |
| | gross photosynthesis | < stony corals | < soft corals | 9 | 9 | 2.451 | 8 | **0.040** |
| | calcification | < stony corals | < soft corals | 9 | 9 | 5.378 | 8 | **<.001** |
| *Por & Scl* | net photosynthesis | < stony corals | < soft corals | 9 | 9 | 3.351 | 8 | **0.010** |
| | respiration | < stony corals | < soft corals | 9 | 9 | 3.177 | 8 | **0.013** |
| | gross photosynthesis | < stony corals | < soft corals | 9 | 9 | 3.424 | 8 | **0.009** |
| | calcification | < stony corals | < soft corals | 9 | 9 | 3.342 | 8 | **0.01** |
| *Mon & Cau* | net photosynthesis | < stony corals | < macro-algae | 9 | 9 | -5.849 | 8 | **<.001** |
| | respiration | < stony corals | < macro-algae | 9 | 9 | -2.602 | 8 | **0.032** |
| | gross photosynthesis | < stony corals | < macro-algae | 9 | 9 | -5.660 | 8 | **<.001** |
| | calcification | < stony corals | < macro-algae | 9 | 9 | 1.544 | 8 | 0.161 |
| *Por & Cau* | net photosynthesis | < stony corals | < macro-algae | 9 | 9 | -2.704 | 8 | **0.027** |
| | respiration | < stony corals | < macro-algae | 9 | 9 | 2.437 | 8 | **0.041** |

| Phase Shift incu-bations | Variable | Group1 | Group2 | n1 | n2 | statistic | df | p.adj |
|---|---|---|---|---|---|---|---|---|
| | gross photosynthesis | < stony corals | < macro-algae | 9 | 9 | -1.587 | 8 | 0.151 |
| | calcification | < stony corals | < macro-algae | 8 | 9 | 0.4 | 7 | 0.701 |
| *Mon & Pey* | net photosynthesis | < stony corals | < macro-algae | 9 | 9 | 3.641 | 8 | **0.007** |
| | respiration | < stony corals | < macro-algae | 9 | 9 | 2.795 | 8 | **0.023** |
| | gross photosynthesis | < stony corals | < macro-algae | 9 | 9 | 3.675 | 8 | **0.006** |
| | calcification | < stony corals | < macro-algae | 9 | 9 | 2.85 | 8 | **0.021** |
| *Por & Pey* | net photosynthesis | < stony corals | < macro-algae | 9 | 9 | 4.618 | 8 | **0.002** |
| | respiration | < stony corals | < macro-algae | 9 | 9 | 3.187 | 8 | **0.013** |
| | gross photosynthesis | < stony corals | < macro-algae | 9 | 9 | 4.342 | 8 | **0.002** |
| | calcification | < stony corals | < macro-algae | 8 | 9 | 0.288 | 7 | 0.782 |

**Tab. S5 Paired t-tests with Bonferroni correction were used to compare measured community productivity in the phase shift incubations with the expected productivity derived from monoculture values of the organisms composing each polyculture (Fig. 4 e-h).** Wilcoxon signed rank tests were applied when parametric assumptions were not met. Significant changes ($p < 0.05$) are indicated in bold

| Phase Shift incubations | Variable | Group1 | Group2 | n1 | n2 | test | statistic | df | p.adj |
|---|---|---|---|---|---|---|---|---|---|
| *Mon & Mon & Xen* | net photosynthesis | measured | expected | 9 | 9 | wilcox | 4 | - | **0.027** |
| | respiration | measured | expected | 9 | 9 | wilcox | 20 | - | 0.82 |
| | gross photosynthesis | measured | expected | 9 | 9 | t - test | -1.771 | 8 | 0.114 |
| | calcification | measured | expected | 9 | 9 | t - test | -0.607 | 8 | 0.561 |
| *Por & Por & Xen* | net photosynthesis | measured | expected | 9 | 9 | wilcox | 8 | - | 0.098 |
| | respiration | measured | expected | 9 | 9 | wilcox | 7 | - | 0.074 |
| | gross photosynthesis | measured | expected | 9 | 9 | t - test | -3.480 | 8 | **0.008** |
| | calcification | measured | expected | 9 | 8 | t - test | -1.910 | 8 | 0.092 |
| *Mon & Mon & Scl* | net photosynthesis | measured | expected | 9 | 9 | wilcox | 3 | - | **0.02** |
| | respiration | measured | expected | 9 | 9 | wilcox | 14 | - | 0.36 |
| | gross photosynthesis | measured | expected | 9 | 9 | t - test | -3.805 | 8 | **0.005** |
| | calcification | measured | expected | 9 | 9 | t - test | -2.281 | 8 | 0.052 |
| *Por & Por & Scl* | net photosynthesis | measured | expected | 9 | 9 | wilcox | 23 | - | 1.0 |
| | respiration | measured | expected | 9 | 9 | wilcox | 22 | - | 1.0 |
| | gross photosynthesis | measured | expected | 9 | 9 | t - test | 0.180 | 8 | 0.861 |
| | calcification | measured | expected | 9 | 9 | t - test | -3.042 | 8 | **0.016** |
| *Mon & Xen & Xen* | net photosynthesis | measured | expected | 9 | 9 | wilcox | 0 | - | **0.004** |
| | respiration | measured | expected | 9 | 9 | wilcox | 0 | - | **0.004** |
| | gross photosynthesis | measured | expected | 9 | 9 | t - test | -9.005 | 8 | **<.001** |
| | calcification | measured | expected | 9 | 9 | t - test | -0.937 | 8 | 0.376 |
| *Por & Xen & Xen* | net photosynthesis | measured | expected | 9 | 9 | wilcox | 0 | - | **0.004** |
| | respiration | measured | expected | 9 | 9 | wilcox | 2 | - | **0.012** |
| | gross photosynthesis | measured | expected | 9 | 9 | t - test | -5.802 | 8 | **<.001** |
| | calcification | measured | expected | 8 | 9 | t - test | -1.494 | 7 | 0.179 |
| *Mon & Scl & Scl* | net photosynthesis | measured | expected | 9 | 9 | wilcox | 0 | - | **0.004** |
| | respiration | measured | expected | 9 | 9 | wilcox | 0 | - | **0.004** |

| Phase Shift incubations | Variable | Group1 | Group2 | n1 | n2 | test | statistic | df | p.adj |
|---|---|---|---|---|---|---|---|---|---|
| | gross photosynthesis | measured | expected | 9 | 9 | t - test | -8.570 | 8 | **<.001** |
| | calcification | measured | expected | 9 | 9 | t - test | -2.157 | 8 | 0.063 |
| *Por & Scl & Scl* | net photosynthesis | measured | expected | 9 | 9 | wilcox | 0 | - | **0.004** |
| | respiration | measured | expected | 9 | 9 | wilcox | 0 | - | **0.004** |
| | gross photosynthesis | measured | expected | 9 | 9 | t - test | -10.47 | 8 | **<.001** |
| | calcification | measured | expected | 9 | 9 | t - test | -1.120 | 8 | 0.295 |
| *Mon & Mon & Cau* | net photosynthesis | measured | expected | 9 | 9 | wilcox | 1 | - | **0.008** |
| | respiration | measured | expected | 9 | 9 | wilcox | 17 | - | 0.57 |
| | gross photosynthesis | measured | expected | 9 | 9 | t - test | -3.176 | 8 | **0.013** |
| | calcification | measured | expected | 9 | 9 | t - test | -1.355 | 8 | 0.212 |
| *Por & Por & Cau* | net photosynthesis | measured | expected | 9 | 9 | wilcox | 8 | - | 0.1 |
| | respiration | measured | expected | 9 | 9 | wilcox | 39 | - | 0.055 |
| | gross photosynthesis | measured | expected | 9 | 9 | t - test | -1.081 | 8 | 0.311 |
| | calcification | measured | expected | 8 | 9 | t - test | -0.045 | 7 | 0.965 |
| *Mon & Mon & Pey* | net photosynthesis | measured | expected | 9 | 9 | wilcox | 10 | - | 0.164 |
| | respiration | measured | expected | 9 | 9 | wilcox | 31 | - | 0.36 |
| | gross photosynthesis | measured | expected | 9 | 9 | t - test | -0.030 | 8 | 0.977 |
| | calcification | measured | expected | 9 | 9 | t - test | -0.495 | 8 | 0.634 |
| *Por & Por & Pey* | net photosynthesis | measured | expected | 9 | 9 | wilcox | 7 | - | 0.074 |
| | respiration | measured | expected | 9 | 9 | wilcox | 24 | - | 0.91 |
| | gross photosynthesis | measured | expected | 9 | 9 | t - test | -1.332 | 8 | 0.22 |
| | calcification | measured | expected | 8 | 9 | t - test | -2.053 | 7 | 0.079 |
| *Mon & Cau & Cau* | net photosynthesis | measured | expected | 9 | 9 | wilcox | 2 | - | **0.012** |
| | respiration | measured | expected | 9 | 9 | wilcox | 18 | - | 0.652 |
| | gross photosynthesis | measured | expected | 9 | 9 | t - test | -2.836 | 8 | **0.02** |
| | calcification | measured | expected | 9 | 9 | t - test | -2.210 | 8 | 0.058 |
| *Por & Cau & Cau* | net photosynthesis | measured | expected | 9 | 9 | wilcox | 0 | - | **0.004** |
| | respiration | measured | expected | 9 | 9 | wilcox | 1 | - | **0.008** |
| | gross photosynthesis | measured | expected | 9 | 9 | t - test | -8.565 | 8 | **<.001** |
| | calcification | measured | expected | 8 | 9 | t - test | -0.917 | 8 | 0.386 |

| Phase Shift incubations | Variable | Group1 | Group2 | n1 | n2 | test | statistic | df | p.adj |
|---|---|---|---|---|---|---|---|---|---|
| *Mon & Pey & Pey* | net photosynthesis | measured | expected | 9 | 9 | wilcox | 0 | - | **0.004** |
| | respiration | measured | expected | 9 | 9 | wilcox | 33 | - | 0.25 |
| | gross photosynthesis | measured | expected | 9 | 9 | t - test | -1.213 | 8 | 0.26 |
| | calcification | measured | expected | 9 | 9 | t - test | -0.741 | 8 | 0.480 |
| *Por & Pey & Pey* | net photosynthesis | measured | expected | 9 | 9 | wilcox | 5 | - | **0.04** |
| | respiration | measured | expected | 9 | 9 | wilcox | 35 | - | 0.164 |
| | gross photosynthesis | measured | expected | 9 | 9 | t - test | 0.353 | 8 | 0.733 |
| | calcification | measured | expected | 8 | 9 | t - test | 0.708 | 8 | 0.499 |

**Tab. S6 One way ANOVA of the 'degraded reef' polyculture assemblages, followed by a Tukey post hoc test with Bonferroni correction, comparing the productivity of the same soft coral and macroalgae species assemblage with and without the two stony coral species (Fig. 5 a-c).** Significant changes ($p < 0.05$) are indicated in bold. For incubations consisting of *S. wanannense* and *C. brachypus*, both respiration and gross photosynthesis rates were log-transformed. For incubations with *X. umbellata* and Peyssonnelia sp., the gross photosynthesis variable was likewise log transformed.

| | | | One-way Anova | | | | | Tukey with Bonferroni correction | | | | |
|---|---|---|---|---|---|---|---|---|---|---|---|---|
| Variable | Combi | | #DF | Sum Sq | Mean Sq | F value | p-value | Groups | diff | lwr | upr | p adj. |
| net photo-synthesis | Pey & Xen | Group | 2 | 0.95 | 0.48 | 0.87 | 0.434 | - | - | - | - | - |
| | | Resid-uals | 24 | 13.2 | 0.55 | | | | | | | |
| | Pey & Scl | Group | 2 | 0.30 | 0.15 | 0.74 | 0.489 | - | - | - | - | - |
| | | Resid-uals | 24 | 4.93 | 0.21 | | | | | | | |
| | Cau & Xen | Group | 2 | 16.9 | 8.45 | 2.02 | 0.155 | - | - | - | - | - |
| | | Resid-uals | 24 | 100 | 4.19 | | | | | | | |
| | Cau & Scl | Group | 2 | 14.6 | 7.28 | 3.47 | **0.048** | Cau-Scl-Por & Cau-Scl-Mon | 0.32 | -1.39 | 2.03 | 0.887 |
| | | Resid-uals | 24 | 50.4 | 2.10 | | | | | | | |
| | | | | | | | | Cau-Scl & Cau-Scl-Mon | -1.37 | -3.08 | 0.33 | 0.13 |
| | | | | | | | | Cau-Scl & Cau-Scl-Por | -1.69 | -3.40 | 0.01 | 0.0521 |

| | | | One-way Anova | | | | | Tukey with Bonferroni correction | | | | |
|---|---|---|---|---|---|---|---|---|---|---|---|---|
| Variable | Combi | | #DF | Sum Sq | Mean Sq | F value | p-value | Groups | diff | lwr | upr | p adj. |
| resp-iration | Pey & Xen | Group<br>Resid-uals | 2<br>22 | 0.45<br>2.76 | 0.23<br>0.13 | 1.79 | 0.190 | - | - | - | - | - |
| | Pey & Scl | Group<br>Resid-uals | 2<br>24 | 0.39<br>2.11 | 0.20<br>0.09 | 2.24 | 0.128 | - | - | - | - | **-** |
| | Cau & Xen | Group<br>Resid-uals | 2<br>24 | 30.4<br>22.2 | 15.2<br>0.92 | 16.4 | **<.001** | Cau-Xen-Por & Cau-Xen-Mon | 0.93 | -0.20 | 2.06 | 0.122 |
| | | | | | | | | Cau-Xen & Cau-Xen-Mon | -1.64 | -2.77 | -0.50 | **0.004** |
| | | | | | | | | Cau-Xen & Cau-Xen-Por | -2.57 | -3.70 | -1.43 | **<.001** |
| | Cau & Scl | Group<br>Resid-uals | 2<br>24 | 2.85<br>3.06 | 1.43<br>0.13 | 11.2 | **<.001** | Cau-Scl-Por & Cau-Scl-Mon | 0.19 | -0.23 | 0.61 | 0.521 |
| | | | | | | | | Cau-Scl & Cau-Scl-Mon | -0.58 | -1.00 | -0.16 | **0.006** |

| | | | One-way Anova | | | | | Tukey with Bonferroni correction | | | | |
|---|---|---|---|---|---|---|---|---|---|---|---|---|
| Variable | Combi | | #DF | Sum Sq | Mean Sq | F value | p-value | Groups | diff | lwr | upr | p adj. |
| | | | | | | | | Cau-Scl & Cau-Scl-Por | -0.76 | -1.18 | -0.34 | **<.001** |
| gross photo-synthesis | Pey & Xen | Group<br>Resid-uals | 2<br>22 | 0.16<br>2.38 | 0.08<br>0.11 | 0.75 | 0.484 | - | - | - | - | - |
| | Pey & Scl | Group<br>Resid-uals | 2<br>24 | 1.28<br>11.3 | 0.64<br>0.47 | 1.36 | 0.277 | - | - | - | - | - |
| | Cau & Xen | Group<br>Resid-uals | 2<br>24 | 77.5<br>131 | 38.7<br>5.46 | 7.09 | **0.004** | Cau-Xen-Por & Cau-Xen-Mon | 2.74 | -0.01 | 5.49 | 0.051 |
| | | | | | | | | Cau-Xen & Cau-Xen-Mon | -1.33 | -4.08 | 1.43 | 0.463 |
| | | | | | | | | Cau-Xen & Cau-Xen-Por | -4.07 | -6.82 | -1.32 | **0.003** |
| | Cau & Scl | Group<br>Resid-uals | 2<br>24 | 1.16<br>2.13 | 0.58<br>0.09 | 6.54 | **0.005** | Cau-Scl-Por & Cau-Scl-Mon | 0.10 | -0.25 | 0.45 | 0.759 |

| | | | One-way Anova | | | | | Tukey with Bonferroni correction | | | | |
|---|---|---|---|---|---|---|---|---|---|---|---|---|
| Variable | Combi | | #DF | Sum Sq | Mean Sq | F value | p-value | Groups | diff | lwr | upr | p adj. |
| | | | | | | | | Cau-Scl & Cau-Scl-Mon | -0.38 | -0.73 | -0.03 | **0.031** |
| | | | | | | | | Cau-Scl & Cau-Scl-Por | -0.48 | -0.83 | -0.13 | **0.006** |

**Tab. S7 Wilcoxen signed rank tests were used to compare measured community productivity in the phase shift incubations with the expected productivity derived from monoculture values of the organisms composing each polyculture (Fig. 5d-f).** Significant changes ($p < 0.05$) are indicated in bold

| Phase Shift incubations | Variable | Group1 | Group2 | n1 | n2 | test | statistic | p |
|---|---|---|---|---|---|---|---|---|
| *Pey & Xen & Mon* | net photosynthesis | measured | expected | 9 | 9 | wilcox | 0 | **0.004** |
| | respiration | measured | expected | 9 | 9 | wilcox | 28 | 0.570 |
| | gross photosynthesis | measured | expected | 9 | 9 | wilcox | 2 | **0.012** |
| *Pey & Xen & Por* | net photosynthesis | measured | expected | 9 | 9 | wilcox | 6 | 0.055 |
| | respiration | measured | expected | 9 | 9 | wilcox | 34 | 0.203 |
| | gross photosynthesis | measured | expected | 9 | 9 | wilcox | 7 | 0.074 |
| *Pey & Xen* | net photosynthesis | measured | expected | 9 | 9 | wilcox | 33 | 0.250 |
| | respiration | measured | expected | 7 | 9 | wilcox | 21 | 0.297 |
| | gross photosynthesis | measured | expected | 7 | 9 | wilcox | 19 | 0.469 |
| *Pey & Scl & Mon* | net photosynthesis | measured | expected | 9 | 9 | wilcox | 0 | **0.004** |
| | respiration | measured | expected | 9 | 9 | wilcox | 6 | 0.055 |
| | gross photosynthesis | measured | expected | 9 | 9 | wilcox | 0 | **0.004** |
| *Pey & Scl & Por* | net photosynthesis | measured | expected | 9 | 9 | wilcox | 0 | **0.004** |
| | respiration | measured | expected | 9 | 9 | wilcox | 5 | **0.039** |
| | gross photosynthesis | measured | expected | 9 | 9 | wilcox | 0 | **0.004** |
| *Pey & Scl* | net photosynthesis | measured | expected | 9 | 9 | wilcox | 13 | 0.301 |
| | respiration | measured | expected | 9 | 9 | wilcox | 22 | 1.000 |
| | gross photosynthesis | measured | expected | 9 | 9 | wilcox | 17 | 0.570 |
| *Cau & Xen & Mon* | net photosynthesis | measured | expected | 9 | 9 | wilcox | 1 | **0.008** |
| | respiration | measured | expected | 9 | 9 | wilcox | 17 | 0.570 |
| | gross photosynthesis | measured | expected | 9 | 9 | wilcox | 1 | **0.008** |
| *Cau & Xen & Por* | net photosynthesis | measured | expected | 9 | 9 | wilcox | 1 | **0.008** |
| | respiration | measured | expected | 9 | 9 | wilcox | 28 | 0.570 |
| | gross photosynthesis | measured | expected | 9 | 9 | wilcox | 4 | **0.027** |
| *Cau & Xen* | net photosynthesis | measured | expected | 9 | 9 | wilcox | 5 | **0.039** |
| | respiration | measured | expected | 9 | 9 | wilcox | 1 | **0.008** |
| | gross photosynthesis | measured | expected | 9 | 9 | wilcox | 2 | **0.012** |

| Phase Shift incubations | Variable | Group1 | Group2 | n1 | n2 | test | statistic | p |
|---|---|---|---|---|---|---|---|---|
| *Cau & Scl & Mon* | net photosynthesis | measured | expected | 9 | 9 | wilcox | 0 | **0.004** |
| | respiration | measured | expected | 9 | 9 | wilcox | 7 | 0.074 |
| | gross photosynthesis | measured | expected | 9 | 9 | wilcox | 0 | **0.004** |
| *Cau & Scl & Por* | net photosynthesis | measured | expected | 9 | 9 | wilcox | 0 | **0.004** |
| | respiration | measured | expected | 9 | 9 | wilcox | 21 | 0.910 |
| | gross photosynthesis | measured | expected | 9 | 9 | wilcox | 0 | **0.004** |
| *Cau & Scl* | net photosynthesis | measured | expected | 9 | 9 | wilcox | 1 | **0.008** |
| | respiration | measured | expected | 9 | 9 | wilcox | 1 | **0.008** |
| | gross photosynthesis | measured | expected | 9 | 9 | wilcox | 0 | **0.004** |

**Tab. S8 One way ANOVA comparing individual photosynthetic efficiency (YII) measurements between monoculture incubations and corresponding polyculture incubations, including coral fragment ID as random factor,** followed by a Tukey post hoc test with Bonferroni correction (Fig. 6). Significant changes ($p < 0.05$) are indicated in bold. Net photosynthesis and gross photosynthesis were log-transformed. Three photosynthetic values, six calcification values and two PAM values were found to be outliers and removed from the dataset.

| | One-way Anova | | | | | | Tukey with Bonferroni correction | | | | |
|---|---|---|---|---|---|---|---|---|---|---|---|
| mea-sured species | | # DF | den DF | F value | p-value | Groups | estimate | Std. error | df | t.ratio | p adj. |
| M. digitata | Int.<br>Mix | 1<br>8 | 112<br>112 | 205600<br>5.66 | **<.001**<br>**<.001** | Mon - Mon_Xen | 0.02 | 0.00 | 112 | 3.71 | **0.010** |
| | | | | | | Mon - Mon_Scl | 0.01 | 0.00 | 112 | 2.07 | 0.501 |
| | | | | | | Mon - Cau_Mon | 0.01 | 0.00 | 112 | 2.94 | 0.089 |
| | | | | | | Mon - Mon_Pey | 0.01 | 0.00 | 112 | 2.07 | 0.501 |
| | | | | | | Mon - Cau_Mon_Xen | 0.00 | 0.00 | 112 | -0.76 | 0.998 |
| | | | | | | Mon - Mon_Pey_Xen | 0.01 | 0.00 | 112 | 1.57 | 0.819 |
| | | | | | | Mon - Cau_Mon_Scl | -0.01 | 0.00 | 112 | -1.48 | 0.862 |
| | | | | | | Mon - Mon_Pey_Scl | -0.01 | 0.00 | 112 | -1.96 | 0.575 |
| | | | | | | Mon_Xen - Mon_Scl | -0.01 | 0.01 | 112 | -1.34 | 0.917 |
| | | | | | | Mon_Xen - Cau_Mon | 0.00 | 0.01 | 112 | -0.62 | 0.999 |
| | | | | | | Mon_Xen - Mon_Pey | -0.01 | 0.01 | 112 | -1.34 | 0.917 |
| | | | | | | Mon_Xen - Cau_Mon_Xen | -0.02 | 0.01 | 112 | -3.64 | **0.012** |
| | | | | | | Mon_Xen - Mon_Pey_Xen | -0.01 | 0.01 | 112 | -1.74 | 0.718 |
| | | | | | | Mon_Xen - Cau_Mon_Scl | -0.02 | 0.01 | 112 | -4.24 | **0.002** |
| | | | | | | Mon_Xen - Mon_Pey_Scl | -0.02 | 0.01 | 112 | -4.63 | **0.000** |
| | | | | | | Mon_Scl - Cau_Mon | 0.00 | 0.01 | 112 | 0.72 | 0.998 |

| | One-way Anova | | | | | | | Tukey with Bonferroni correction | | | |
|---|---|---|---|---|---|---|---|---|---|---|---|
| mea-sured species | | # DF | den DF | F value | p-value | Groups | estimate | Std. error | df | t.ratio | p adj. |
| | | | | | | Mon_Scl - Mon_Pey | 0.00 | 0.01 | 112 | 0.00 | 1.000 |
| | | | | | | Mon_Scl - Cau_Mon_Xen | -0.01 | 0.01 | 112 | -2.30 | 0.348 |
| | | | | | | Mon_Scl - Mon_Pey_Xen | 0.00 | 0.01 | 112 | -0.40 | 1.000 |
| | | | | | | Mon_Scl - Cau_Mon_Scl | -0.02 | 0.01 | 112 | -2.90 | 0.101 |
| | | | | | | Mon_Scl - Mon_Pey_Scl | -0.02 | 0.01 | 112 | -3.29 | **0.035** |
| | | | | | | Cau_Mon - Mon_Pey | 0.00 | 0.01 | 112 | -0.72 | 0.998 |
| | | | | | | Cau_Mon - Cau_Mon_Xen | -0.02 | 0.01 | 112 | -3.02 | 0.073 |
| | | | | | | Cau_Mon - Mon_Pey_Xen | -0.01 | 0.01 | 112 | -1.12 | 0.970 |
| | | | | | | Cau_Mon - Cau_Mon_Scl | -0.02 | 0.01 | 112 | -3.61 | **0.013** |
| | | | | | | Cau_Mon - Mon_Pey_Scl | -0.02 | 0.01 | 112 | -4.00 | **0.003** |
| | | | | | | Mon_Pey - Cau_Mon_Xen | -0.01 | 0.01 | 112 | -2.30 | 0.348 |
| | | | | | | Mon_Pey - Mon_Pey_Xen | 0.00 | 0.01 | 112 | -0.40 | 1.000 |
| | | | | | | Mon_Pey - Cau_Mon_Scl | -0.02 | 0.01 | 112 | -2.90 | 0.101 |
| | | | | | | Mon_Pey - Mon_Pey_Scl | -0.02 | 0.01 | 112 | -3.29 | **0.035** |
| | | | | | | Cau_Mon_Xen - Mon_Pey_Xen | 0.01 | 0.01 | 112 | 1.90 | 0.615 |
| | | | | | | Cau_Mon_Xen - Cau_Mon_Scl | 0.00 | 0.01 | 112 | -0.59 | 1.000 |
| | | | | | | Cau_Mon_Xen - Mon_Pey_Scl | -0.01 | 0.01 | 112 | -0.98 | 0.987 |
| | | | | | | Mon_Pey_Xen - Cau_Mon_Scl | -0.01 | 0.01 | 112 | -2.49 | 0.247 |

| | One-way Anova | | | | | | Tukey with Bonferroni correction | | | | |
|---|---|---|---|---|---|---|---|---|---|---|---|
| mea-sured species | | # DF | den DF | F value | p-value | Groups | estimate | Std. error | df | t.ratio | p adj. |
| | | | | | | Mon_Pey_Xen - Mon_Pey_Scl | -0.02 | 0.01 | 112 | -2.88 | 0.105 |
| | | | | | | Cau_Mon_Scl - Mon_Pey_Scl | 0.00 | 0.01 | 112 | -0.39 | 1.000 |
| P. rus | Int. | 1 | 111 | 6401 | **<.001** | Por -Por_Xen | 0.01 | 0.01 | 111 | 2.00 | 0.547 |
| | Mix | 8 | 111 | 2.55 | **0.014** | Por - Por_Scl | 0.02 | 0.01 | 111 | 2.82 | 0.120 |
| | | | | | | Por - Cau_Por | 0.00 | 0.01 | 111 | -0.17 | 1.000 |
| | | | | | | Por - Pey_Por | 0.00 | 0.01 | 111 | -0.15 | 1.000 |
| | | | | | | Por - Cau_Por_Xen | -0.01 | 0.01 | 111 | -1.85 | 0.650 |
| | | | | | | Por - Pey_Por_Xen | 0.00 | 0.01 | 111 | -0.10 | 1.000 |
| | | | | | | Por - Cau_Por_Scl | 0.00 | 0.01 | 111 | -0.32 | 1.000 |
| | | | | | | Por - Pey_Por_Scl | 0.00 | 0.01 | 111 | -0.29 | 1.000 |
| | | | | | | Por_Xen - Por_Scl | 0.01 | 0.01 | 111 | 0.67 | 0.999 |
| | | | | | | Por_Xen - Cau_Por | -0.01 | 0.01 | 111 | -1.77 | 0.698 |
| | | | | | | Por_Xen - Pey_Por | -0.01 | 0.01 | 111 | -1.75 | 0.711 |
| | | | | | | Por_Xen - Cau_Por_Xen | -0.03 | 0.01 | 111 | -3.14 | 0.053 |
| | | | | | | Por_Xen - Pey_Por_Xen | -0.01 | 0.01 | 111 | -1.71 | 0.737 |
| | | | | | | Por_Xen - Cau_Por_Scl | -0.02 | 0.01 | 111 | -1.90 | 0.617 |
| | | | | | | Por_Xen - Pey_Por_Scl | -0.02 | 0.01 | 111 | -1.83 | 0.660 |
| | | | | | | Por_Scl - Cau_Por | -0.02 | 0.01 | 111 | -2.45 | 0.269 |
| | | | | | | Por_Scl - Pey_Por | -0.02 | 0.01 | 111 | -2.43 | 0.280 |

| | One-way Anova | | | | | | Tukey with Bonferroni correction | | | | |
|---|---|---|---|---|---|---|---|---|---|---|---|
| mea-sured species | | # DF | den DF | F value | p-value | Groups | estimate | Std. error | df | t.ratio | p adj. |
| | | | | | | Por_Scl - Cau_Por_Xen | -0.03 | 0.01 | 111 | -3.82 | **0.007** |
| | | | | | | Por_Scl - Pey_Por_Xen | -0.02 | 0.01 | 111 | -2.39 | 0.301 |
| | | | | | | Por_Scl - Cau_Por_Scl | -0.02 | 0.01 | 111 | -2.57 | 0.211 |
| | | | | | | Por_Scl - Pey_Por_Scl | -0.02 | 0.01 | 111 | -2.49 | 0.248 |
| | | | | | | Cau_Por - Pey_Por | 0.00 | 0.01 | 111 | 0.02 | 1.000 |
| | | | | | | Cau_Por - Cau_Por_Xen | -0.01 | 0.01 | 111 | -1.37 | 0.908 |
| | | | | | | Cau_Por - Pey_Por_Xen | 0.00 | 0.01 | 111 | 0.06 | 1.000 |
| | | | | | | Cau_Por - Cau_Por_Scl | 0.00 | 0.01 | 111 | -0.12 | 1.000 |
| | | | | | | Cau_Por - Pey_Por_Scl | 0.00 | 0.01 | 111 | -0.10 | 1.000 |
| | | | | | | Pey_Por - Cau_Por_Xen | -0.01 | 0.01 | 111 | -1.39 | 0.900 |
| | | | | | | Pey_Por - Pey_Por_Xen | 0.00 | 0.01 | 111 | 0.04 | 1.000 |
| | | | | | | Pey_Por - Cau_Por_Scl | 0.00 | 0.01 | 111 | -0.14 | 1.000 |
| | | | | | | Pey_Por - Pey_Por_Scl | 0.00 | 0.01 | 111 | -0.12 | 1.000 |
| | | | | | | Cau_Por_Xen - Pey_Por_Xen | 0.01 | 0.01 | 111 | 1.43 | 0.885 |
| | | | | | | Cau_Por_Xen - Cau_Por_Scl | 0.01 | 0.01 | 111 | 1.24 | 0.945 |
| | | | | | | Cau_Por_Xen - Pey_Por_Scl | 0.01 | 0.01 | 111 | 1.23 | 0.947 |
| | | | | | | Pey_Por_Xen - Cau_Por_Scl | 0.00 | 0.01 | 111 | -0.18 | 1.000 |
| | | | | | | Pey_Por_Xen - Pey_Por_Scl | 0.00 | 0.01 | 111 | -0.16 | 1.000 |

| | One-way Anova | | | | | | Tukey with Bonferroni correction | | | | |
|---|---|---|---|---|---|---|---|---|---|---|---|
| mea-sured species | | # DF | den DF | F value | p-value | Groups | estimate | Std. error | df | t.ratio | p adj. |
| | | | | | | Cau_Por_Scl - Pey_Por_Scl | 0.00 | 0.01 | 111 | 0.02 | 1.000 |
| Xenia | Int.<br>Mix | 1<br>8 | 111<br>111 | 9153<br>2.07 | <.001 | Xen - Mon_Xen | 0.01 | 0.01 | 111 | 1.11 | 0.971 |
| | | | | | | Xen - Por_Xen | 0.01 | 0.01 | 111 | 0.98 | 0.987 |
| | | | | | | Xen - Cau_Xen | -0.01 | 0.01 | 111 | -1.28 | 0.935 |
| | | | | | | Xen - Pey_Xen | -0.01 | 0.01 | 111 | -1.34 | 0.916 |
| | | | | | | Xen - Cau_Mon_Xen | 0.00 | 0.01 | 111 | 0.28 | 1.000 |
| | | | | | | Xen - Mon_Pey_Xen | 0.00 | 0.01 | 111 | 0.45 | 1.000 |
| | | | | | | Xen - Cau_Por_Xen | -0.02 | 0.01 | 111 | -2.55 | 0.220 |
| | | | | | | Xen - Pey_Por_Xen | -0.01 | 0.01 | 111 | -1.02 | 0.983 |
| | | | | | | Mon_Xen - Por_Xen | 0.00 | 0.01 | 111 | -0.11 | 1.000 |
| | | | | | | Mon_Xen - Cau_Xen | -0.02 | 0.01 | 111 | -1.96 | 0.577 |
| | | | | | | Mon_Xen - Pey_Xen | -0.02 | 0.01 | 111 | -2.00 | 0.550 |
| | | | | | | Mon_Xen - Cau_Mon_Xen | -0.01 | 0.01 | 111 | -0.68 | 0.999 |
| | | | | | | Mon_Xen - Mon_Pey_Xen | 0.00 | 0.01 | 111 | -0.54 | 1.000 |
| | | | | | | Mon_Xen - Cau_Por_Xen | -0.03 | 0.01 | 111 | -2.99 | 0.079 |
| | | | | | | Mon_Xen - Pey_Por_Xen | -0.01 | 0.01 | 111 | -1.74 | 0.720 |
| | | | | | | Por_Xen - Cau_Xen | -0.02 | 0.01 | 111 | -1.85 | 0.651 |
| | | | | | | Por_Xen - Pey_Xen | -0.02 | 0.01 | 111 | -1.89 | 0.622 |

| | One-way Anova | | | | | | | | Tukey with Bonferroni correction | | |
|---|---|---|---|---|---|---|---|---|---|---|---|
| mea-sured species | | # DF | den DF | F value | p-value | Groups | estimate | Std. error | df | t.ratio | p adj. |
| | | | | | | Por_Xen - Cau_Mon_Xen | 0.00 | 0.01 | 111 | -0.58 | 1.000 |
| | | | | | | Por_Xen - Mon_Pey_Xen | 0.00 | 0.01 | 111 | -0.43 | 1.000 |
| | | | | | | Por_Xen - Cau_Por_Xen | -0.02 | 0.01 | 111 | -2.88 | 0.104 |
| | | | | | | Por_Xen - Pey_Por_Xen | -0.01 | 0.01 | 111 | -1.63 | 0.785 |
| | | | | | | Cau_Xen - Pey_Xen | 0.00 | 0.01 | 111 | -0.09 | 1.000 |
| | | | | | | Cau_Xen - Cau_Mon_Xen | 0.01 | 0.01 | 111 | 1.27 | 0.938 |
| | | | | | | Cau_Xen - Mon_Pey_Xen | 0.01 | 0.01 | 111 | 1.42 | 0.889 |
| | | | | | | Cau_Xen - Cau_Por_Xen | -0.01 | 0.01 | 111 | -1.04 | 0.981 |
| | | | | | | Cau_Xen - Pey_Por_Xen | 0.00 | 0.01 | 111 | 0.22 | 1.000 |
| | | | | | | Pey_Xen - Cau_Mon_Xen | 0.01 | 0.01 | 111 | 1.33 | 0.921 |
| | | | | | | Pey_Xen - Mon_Pey_Xen | 0.01 | 0.01 | 111 | 1.47 | 0.867 |
| | | | | | | Pey_Xen - Cau_Por_Xen | -0.01 | 0.01 | 111 | -0.92 | 0.991 |
| | | | | | | Pey_Xen - Pey_Por_Xen | 0.00 | 0.01 | 111 | 0.30 | 1.000 |
| | | | | | | Cau_Mon_Xen - Mon_Pey_Xen | 0.00 | 0.01 | 111 | 0.15 | 1.000 |
| | | | | | | Cau_Mon_Xen - Cau_Por_Xen | -0.02 | 0.01 | 111 | -2.31 | 0.347 |
| | | | | | | Cau_Mon_Xen - Pey_Por_Xen | -0.01 | 0.01 | 111 | -1.06 | 0.979 |
| | | | | | | Mon_Pey_Xen - Cau_Por_Xen | -0.02 | 0.01 | 111 | -2.45 | 0.266 |
| | | | | | | Mon_Pey_Xen - Pey_Por_Xen | -0.01 | 0.01 | 111 | -1.20 | 0.954 |

| | One-way Anova | | | | | | Tukey with Bonferroni correction | | | | |
|---|---|---|---|---|---|---|---|---|---|---|---|
| mea-sured species | | # DF | den DF | F value | p-value | Groups | estimate | Std. error | df | t.ratio | p adj. |
| | | | | | | Cau_Por_Xen - Pey_Por_Xen | 0.01 | 0.01 | 111 | 1.25 | 0.943 |
| Scl | Int.<br>Mix | 1<br>8 | 112<br>112 | 7500<br>1.38 | <.001<br>0.211 | Scl - Mon_Scl | 0.01 | 0.01 | 112 | 2.08 | 0.489 |
| | | | | | | Scl - Por_Scl | 0.01 | 0.01 | 112 | 1.53 | 0.840 |
| | | | | | | Scl - Cau_Scl | 0.01 | 0.01 | 112 | 0.82 | 0.996 |
| | | | | | | Scl - Pey_Scl | 0.01 | 0.01 | 112 | 1.18 | 0.959 |
| | | | | | | Scl - Cau_Mon_Scl | 0.00 | 0.01 | 112 | 0.26 | 1.000 |
| | | | | | | Scl - Mon_Pey_Scl | -0.01 | 0.01 | 112 | -1.15 | 0.965 |
| | | | | | | Scl - Cau_Por_Scl | 0.00 | 0.01 | 112 | 0.21 | 1.000 |
| | | | | | | Scl - Pey_Por_Scl | 0.00 | 0.01 | 112 | -0.29 | 1.000 |
| | | | | | | Mon_Scl - Por_Scl | 0.00 | 0.01 | 112 | -0.46 | 1.000 |
| | | | | | | Mon_Scl - Cau_Scl | -0.01 | 0.01 | 112 | -1.03 | 0.982 |
| | | | | | | Mon_Scl - Pey_Scl | -0.01 | 0.01 | 112 | -0.74 | 0.998 |
| | | | | | | Mon_Scl - Cau_Mon_Scl | -0.01 | 0.01 | 112 | -1.49 | 0.859 |
| | | | | | | Mon_Scl - Mon_Pey_Scl | -0.02 | 0.01 | 112 | -2.64 | 0.182 |
| | | | | | | Mon_Scl - Cau_Por_Scl | -0.01 | 0.01 | 112 | -1.53 | 0.840 |
| | | | | | | Mon_Scl - Pey_Por_Scl | -0.02 | 0.01 | 112 | -1.94 | 0.587 |
| | | | | | | Por_Scl - Cau_Scl | 0.00 | 0.01 | 112 | -0.58 | 1.000 |
| | | | | | | Por_Scl - Pey_Scl | 0.00 | 0.01 | 112 | -0.28 | 1.000 |

| | One-way Anova | | | | | | Tukey with Bonferroni correction | | | | |
|---|---|---|---|---|---|---|---|---|---|---|---|
| mea-sured species | | # DF | den DF | F value | p-value | Groups | estimate | Std. error | df | t.ratio | p adj. |
| | | | | | | Por_Scl - Cau_Mon_Scl | -0.01 | 0.01 | 112 | -1.03 | 0.982 |
| | | | | | | Por_Scl - Mon_Pey_Scl | -0.02 | 0.01 | 112 | -2.18 | 0.423 |
| | | | | | | Por_Scl - Cau_Por_Scl | -0.01 | 0.01 | 112 | -1.07 | 0.977 |
| | | | | | | Por_Scl - Pey_Por_Scl | -0.01 | 0.01 | 112 | -1.49 | 0.859 |
| | | | | | | Cau_Scl - Pey_Scl | 0.00 | 0.01 | 112 | 0.29 | 1.000 |
| | | | | | | Cau_Scl - Cau_Mon_Scl | 0.00 | 0.01 | 112 | -0.46 | 1.000 |
| | | | | | | Cau_Scl - Mon_Pey_Scl | -0.01 | 0.01 | 112 | -1.61 | 0.799 |
| | | | | | | Cau_Scl - Cau_Por_Scl | 0.00 | 0.01 | 112 | -0.50 | 1.000 |
| | | | | | | Cau_Scl - Pey_Por_Scl | -0.01 | 0.01 | 112 | -0.91 | 0.992 |
| | | | | | | Pey_Scl - Cau_Mon_Scl | -0.01 | 0.01 | 112 | -0.75 | 0.998 |
| | | | | | | Pey_Scl - Mon_Pey_Scl | -0.02 | 0.01 | 112 | -1.90 | 0.614 |
| | | | | | | Pey_Scl - Cau_Por_Scl | -0.01 | 0.01 | 112 | -0.79 | 0.997 |
| | | | | | | Pey_Scl - Pey_Por_Scl | -0.01 | 0.01 | 112 | -1.20 | 0.954 |
| | | | | | | Cau_Mon_Scl - Mon_Pey_Scl | -0.01 | 0.01 | 112 | -1.15 | 0.964 |
| | | | | | | Cau_Mon_Scl - Cau_Por_Scl | 0.00 | 0.01 | 112 | -0.04 | 1.000 |
| | | | | | | Cau_Mon_Scl - Pey_Por_Scl | 0.00 | 0.01 | 112 | -0.46 | 1.000 |
| | | | | | | Mon_Pey_Scl - Cau_Por_Scl | 0.01 | 0.01 | 112 | 1.11 | 0.971 |
| | | | | | | Mon_Pey_Scl - Pey_Por_Scl | 0.01 | 0.01 | 112 | 0.70 | 0.999 |

| | One-way Anova | | | | | | Tukey with Bonferroni correction | | | | |
|---|---|---|---|---|---|---|---|---|---|---|---|
| mea-sured species | | # DF | den DF | F value | p-value | Groups | estimate | Std. error | df | t.ratio | p adj. |
| | | | | | | Cau_Por_Scl - Pey_Por_Scl | 0.00 | 0.01 | 112 | -0.41 | 1.000 |
| Pey | Int.<br>Mix | 1<br>8 | 110<br>110 | 5866<br>6.25 | <.001<br><.001 | Pey - Mon_Pey | -0.03 | 0.01 | 110 | -4.12 | **0.002** |
| | | | | | | Pey - Pey_Por | -0.04 | 0.01 | 110 | -5.46 | **0.000** |
| | | | | | | Pey - Pey_Xen | -0.04 | 0.01 | 110 | -4.52 | **0.001** |
| | | | | | | Pey - Pey_Scl | -0.03 | 0.01 | 110 | -4.05 | **0.003** |
| | | | | | | Pey - Mon_Pey_Xen | -0.02 | 0.01 | 110 | -1.97 | 0.565 |
| | | | | | | Pey - Mon_Pey_Scl | -0.02 | 0.01 | 110 | -2.10 | 0.477 |
| | | | | | | Pey - Pey_Por_Xen | -0.01 | 0.01 | 110 | -1.83 | 0.660 |
| | | | | | | Pey - Pey_Por_Scl | -0.01 | 0.01 | 110 | -1.65 | 0.775 |
| | | | | | | Mon_Pey - Pey_Por | -0.01 | 0.01 | 110 | -1.10 | 0.974 |
| | | | | | | Mon_Pey - Pey_Xen | 0.00 | 0.01 | 110 | -0.33 | 1.000 |
| | | | | | | Mon_Pey - Pey_Scl | 0.00 | 0.01 | 110 | 0.06 | 1.000 |
| | | | | | | Mon_Pey - Mon_Pey_Xen | 0.02 | 0.01 | 110 | 1.77 | 0.702 |
| | | | | | | Mon_Pey - Mon_Pey_Scl | 0.02 | 0.01 | 110 | 1.66 | 0.767 |
| | | | | | | Mon_Pey - Pey_Por_Xen | 0.02 | 0.01 | 110 | 1.88 | 0.626 |
| | | | | | | Mon_Pey - Pey_Por_Scl | 0.02 | 0.01 | 110 | 2.03 | 0.523 |
| | | | | | | Pey_Por - Pey_Xen | 0.01 | 0.01 | 110 | 0.77 | 0.997 |
| | | | | | | Pey_Por - Pey_Scl | 0.01 | 0.01 | 110 | 1.16 | 0.963 |

| | One-way Anova | | | | | | Tukey with Bonferroni correction | | | | |
|---|---|---|---|---|---|---|---|---|---|---|---|
| mea-sured species | | # DF | den DF | F value | p-value | Groups | estimate | Std. error | df | t.ratio | p adj. |
| | | | | | | Pey_Por - Mon_Pey_Xen | 0.03 | 0.01 | 110 | 2.87 | 0.109 |
| | | | | | | Pey_Por - Mon_Pey_Scl | 0.03 | 0.01 | 110 | 2.76 | 0.140 |
| | | | | | | Pey_Por - Pey_Por_Xen | 0.03 | 0.01 | 110 | 2.98 | 0.081 |
| | | | | | | Pey_Por - Pey_Por_Scl | 0.03 | 0.01 | 110 | 3.13 | 0.055 |
| | | | | | | Pey_Xen - Pey_Scl | 0.00 | 0.01 | 110 | 0.39 | 1.000 |
| | | | | | | Pey_Xen - Mon_Pey_Xen | 0.02 | 0.01 | 110 | 2.10 | 0.481 |
| | | | | | | Pey_Xen - Mon_Pey_Scl | 0.02 | 0.01 | 110 | 1.99 | 0.553 |
| | | | | | | Pey_Xen - Pey_Por_Xen | 0.02 | 0.01 | 110 | 2.21 | 0.406 |
| | | | | | | Pey_Xen - Pey_Por_Scl | 0.02 | 0.01 | 110 | 2.36 | 0.315 |
| | | | | | | Pey_Scl - Mon_Pey_Xen | 0.02 | 0.01 | 110 | 1.71 | 0.741 |
| | | | | | | Pey_Scl - Mon_Pey_Scl | 0.02 | 0.01 | 110 | 1.60 | 0.802 |
| | | | | | | Pey_Scl - Pey_Por_Xen | 0.02 | 0.01 | 110 | 1.82 | 0.667 |
| | | | | | | Pey_Scl - Pey_Por_Scl | 0.02 | 0.01 | 110 | 1.97 | 0.565 |
| | | | | | | Mon_Pey_Xen - Mon_Pey_Scl | 0.00 | 0.01 | 110 | -0.11 | 1.000 |
| | | | | | | Mon_Pey_Xen - Pey_Por_Xen | 0.00 | 0.01 | 110 | 0.12 | 1.000 |
| | | | | | | Mon_Pey_Xen - Pey_Por_Scl | 0.00 | 0.01 | 110 | 0.27 | 1.000 |
| | | | | | | Mon_Pey_Scl - Pey_Por_Xen | 0.00 | 0.01 | 110 | 0.22 | 1.000 |
| | | | | | | Mon_Pey_Scl - Pey_Por_Scl | 0.00 | 0.01 | 110 | 0.37 | 1.000 |

| | One-way Anova | | | | | | Tukey with Bonferroni correction | | | | |
|---|---|---|---|---|---|---|---|---|---|---|---|
| mea-sured species | | # DF | den DF | F value | p-value | Groups | estimate | Std. error | df | t.ratio | p adj. |
| | | | | | | Pey_Por_Xen - Pey_Por_Scl | 0.00 | 0.01 | 110 | 0.15 | 1.000 |
| Cau | Int.<br>Mix | 1<br>8 | 112<br>112 | 48838<br>6.53 | <.001<br><.001 | Cau - Cau_Mon | 0.03 | 0.01 | 112 | 3.39 | **0.026** |
| | | | | | | Cau - Cau_Por | -0.01 | 0.01 | 112 | -1.46 | 0.872 |
| | | | | | | Cau - Cau_Xen | 0.02 | 0.01 | 112 | 2.50 | 0.241 |
| | | | | | | Cau - Cau_Scl | 0.02 | 0.01 | 112 | 2.50 | 0.246 |
| | | | | | | Cau - Cau_Mon_Xen | -0.01 | 0.01 | 112 | -1.14 | 0.966 |
| | | | | | | Cau - Cau_Mon_Scl | -0.03 | 0.01 | 112 | -2.87 | 0.108 |
| | | | | | | Cau - Cau_Por_Xen | -0.01 | 0.01 | 112 | -1.61 | 0.798 |
| | | | | | | Cau - Cau_Por_Scl | -0.01 | 0.01 | 112 | -0.98 | 0.987 |
| | | | | | | Cau_Mon - Cau_Por | -0.04 | 0.01 | 112 | -3.96 | **0.004** |
| | | | | | | Cau_Mon - Cau_Xen | -0.01 | 0.01 | 112 | -0.72 | 0.998 |
| | | | | | | Cau_Mon - Cau_Scl | -0.01 | 0.01 | 112 | -0.73 | 0.998 |
| | | | | | | Cau_Mon - Cau_Mon_Xen | -0.04 | 0.01 | 112 | -3.70 | **0.010** |
| | | | | | | Cau_Mon - Cau_Mon_Scl | -0.05 | 0.01 | 112 | -5.11 | **0.000** |
| | | | | | | Cau_Mon - Cau_Por_Xen | -0.04 | 0.01 | 112 | -4.08 | **0.003** |
| | | | | | | Cau_Mon - Cau_Por_Scl | -0.04 | 0.01 | 112 | -3.57 | **0.015** |
| | | | | | | Cau_Por - Cau_Xen | 0.03 | 0.01 | 112 | 3.23 | **0.041** |
| | | | | | | Cau_Por - Cau_Scl | 0.03 | 0.01 | 112 | 3.23 | **0.042** |

| | One-way Anova | | | | | | | | | Tukey with Bonferroni correction |
|---|---|---|---|---|---|---|---|---|---|---|
| mea-sured species | | # DF | den DF | F value | p-value | Groups | estimate | Std. error | df | t.ratio | p adj. |
| | | | | | | Cau_Por - Cau_Mon_Xen | 0.00 | 0.01 | 112 | 0.26 | 1.000 |
| | | | | | | Cau_Por - Cau_Mon_Scl | -0.01 | 0.01 | 112 | -1.15 | 0.965 |
| | | | | | | Cau_Por - Cau_Por_Xen | 0.00 | 0.01 | 112 | -0.12 | 1.000 |
| | | | | | | Cau_Por - Cau_Por_Scl | 0.00 | 0.01 | 112 | 0.39 | 1.000 |
| | | | | | | Cau_Xen - Cau_Scl | 0.00 | 0.01 | 112 | -0.01 | 1.000 |
| | | | | | | Cau_Xen - Cau_Mon_Xen | -0.03 | 0.01 | 112 | -2.98 | 0.082 |
| | | | | | | Cau_Xen - Cau_Mon_Scl | -0.05 | 0.01 | 112 | -4.39 | **0.001** |
| | | | | | | Cau_Xen - Cau_Por_Xen | -0.04 | 0.01 | 112 | -3.36 | **0.028** |
| | | | | | | Cau_Xen - Cau_Por_Scl | -0.03 | 0.01 | 112 | -2.85 | 0.114 |
| | | | | | | Cau_Scl - Cau_Mon_Xen | -0.03 | 0.01 | 112 | -2.97 | 0.084 |
| | | | | | | Cau_Scl - Cau_Mon_Scl | -0.05 | 0.01 | 112 | -4.38 | **0.001** |
| | | | | | | Cau_Scl - Cau_Por_Xen | -0.04 | 0.01 | 112 | -3.35 | **0.029** |
| | | | | | | Cau_Scl - Cau_Por_Scl | -0.03 | 0.01 | 112 | -2.84 | 0.116 |
| | | | | | | Cau_Mon_Xen - Cau_Mon_Scl | -0.02 | 0.01 | 112 | -1.41 | 0.893 |
| | | | | | | Cau_Mon_Xen - Cau_Por_Xen | 0.00 | 0.01 | 112 | -0.38 | 1.000 |
| | | | | | | Cau_Mon_Xen - Cau_Por_Scl | 0.00 | 0.01 | 112 | 0.13 | 1.000 |
| | | | | | | Cau_Mon_Scl - Cau_Por_Xen | 0.01 | 0.01 | 112 | 1.03 | 0.983 |
| | | | | | | Cau_Mon_Scl - Cau_Por_Scl | 0.02 | 0.01 | 112 | 1.54 | 0.834 |

| | One-way Anova | | | | | | Tukey with Bonferroni correction | | | | |
|---|---|---|---|---|---|---|---|---|---|---|---|
| mea-sured species | | # DF | den DF | F value | p-value | Groups | estimate | Std. error | df | t.ratio | p adj. |
| | | | | | | Cau_Por_Xen - Cau_Por_Scl | 0.01 | 0.01 | 112 | 0.51 | 1.000 |

### Supplementary Figures

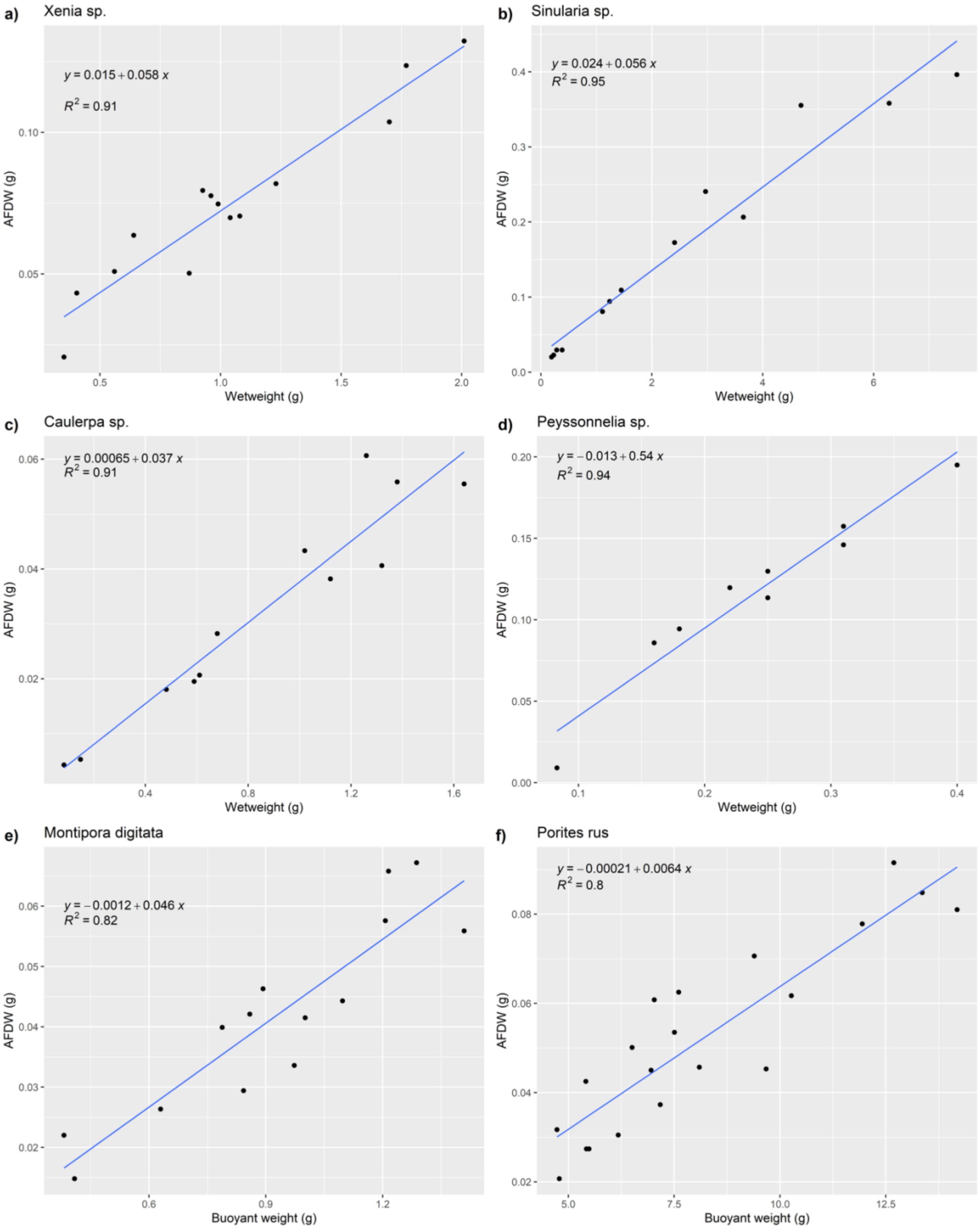

**Figure S1 Correlation between wet weight/ buoyant weight and Ash free dry weight (AFDW) per species**

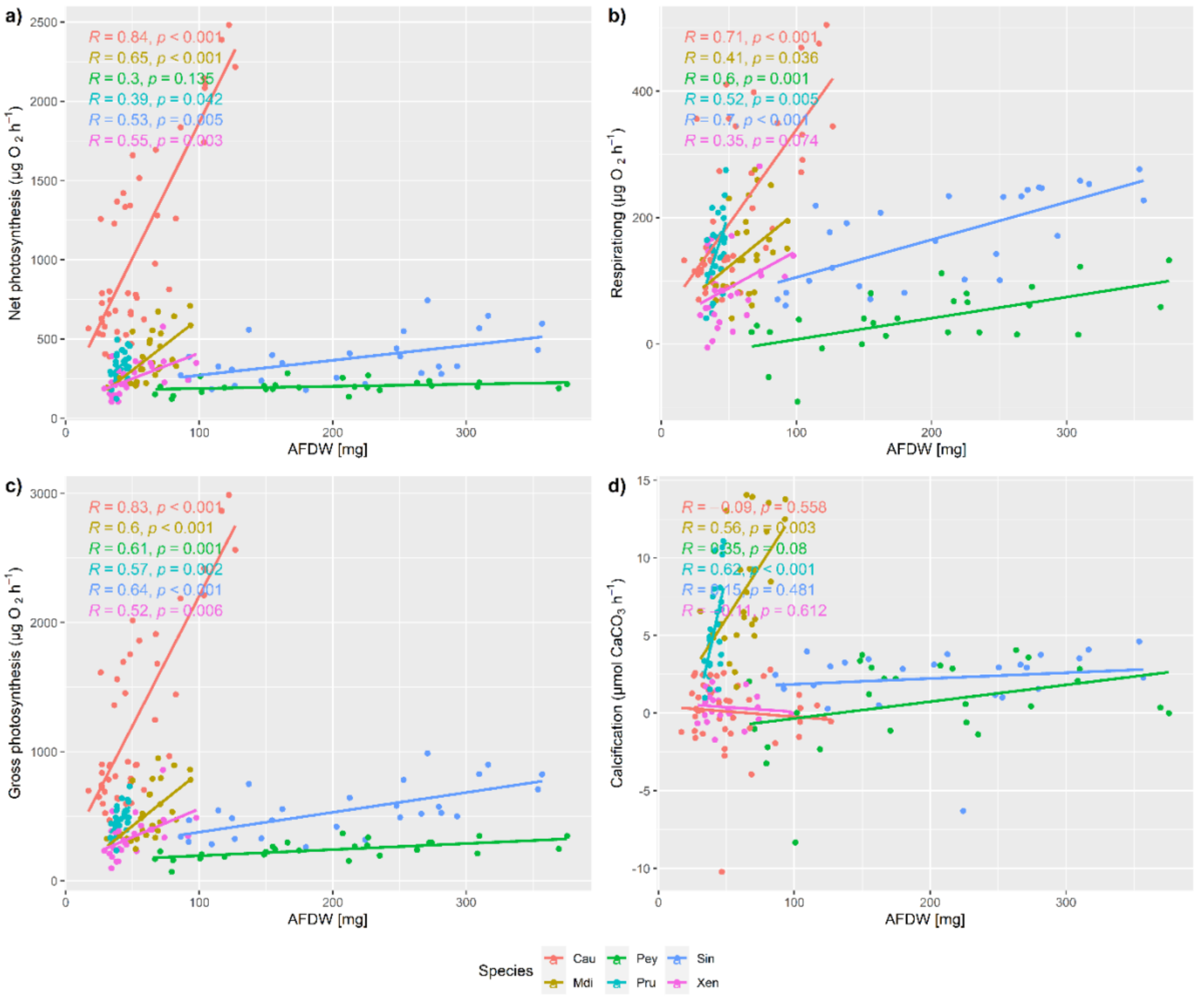

**Figure S2 Pearson correlation between AFDW and productivity parameters** a) net photosynthesis, b) respiration, c) gross photosynthesis, d) calcification
